# The ATPase cycle of Human Muscle Myosin II Isoforms: Adaptation of a single mechanochemical cycle for different physiological roles

**DOI:** 10.1101/672030

**Authors:** Chloe A. Johnson, Jonathan Walklate, Marina Svicevic, Srboljub M. Mijailovich, Carlos Vera, Anastasia Karabina, Leslie A. Leinwand, Michael A. Geeves

## Abstract

Striated muscle myosins are encoded by a large gene family in all mammals, including human. These isoforms define several of the key characteristics of the different striated muscle fiber types including maximum shortening velocity. We have previously used recombinant isoforms of the motor domains of eight different human myosin isoforms to define the actin.myosin cross-bridge cycle in solution. Here, we use a recently developed modeling approach MUSICO to explore how well the experimentally defined cross-bridge cycles for each isoform in solution can predict the characteristics of muscle fiber contraction including duty ratio, shortening velocity, ATP economy and the load dependence of these parameters. The work shows that the parameters of the cross-bridge cycle predict many of the major characteristics of each muscle fiber type and raises the question of what sequence changes are responsible for these characteristics.

## Introduction

Muscle myosin in all mammals consists of a variety of isoforms, each expressed from its own gene (see Table 1 and references therein). There are 10 such genes in the human genome, and one pseudogene. All of the striated muscle myosin sequences are very highly conserved, but each has functional differences and these differences are required for normal muscle function since myosin isoform-specific null mice can have profound phenotypes (1). Further, disease-causing mutations in 6 of the 10 genes have been reported (2, 3). The expression of these genes is regulated temporally and spatially and can be affected by physical activity, animal species and hormonal status. Each myosin isoform confers distinct contractile characteristics to each muscle fiber type (4, 5). These characteristics include maximum shortening velocity, rate of ATP usage, the economy of ATP usage and the velocity at which power output is maximal. Other parameters such as maximal force per cross-bridge or step size are much less variable (4, 5). How each myosin is tuned for its specific function is not well understood. Also poorly understood is how changes in the amino acid sequence of each myosin bring about the functional changes. As a first step to understand how striated muscle myosin has evolved to have distinct physiological roles, we first needed to understand how the ATP driven cross-bridge cycle varies between the isoforms. Towards that end, we recently published a complete characterization of the kinetics of the ATPase cycle for the motor domains of six, adult human muscle-myosin isoforms and the embryonic isoform ((6–8), see Table 1). Here we extend this data set to include the first study of the human perinatal myosin isoform. With this large set of isoform data, it is possible to examine the extent to which differences in the ATPase cycle for each isoform can predict the differences in mechanochemical properties of muscle fibers expressing them. Here we use the recently developed MUSICO modeling approach (9, 10) to predict the contraction characteristics of each muscle fiber type and compare the predictions to published data for single muscle fibers expressing single myosin isoforms.

**Table 1.** Summary of Isoforms.

| ***Isoform*** | ***Human Gene*** | ***Human tissue expression†*** | ***Maximum shortening velocity in muscle fiber\**** | ***Light chains\*\**** | ***Expression tags on heavy chain*** |
| --- | --- | --- | --- | --- | --- |
| alpha-cardiac ($\alpha$) | *MYH6* | Atrial myocardium | | Endogenous Mouse | 6xHis |
| beta-cardiac/slow muscle I ($\beta$) | *MYH7* | Ventricular myocardium/slow skeletal | $0.330 \pm 0.022$ | Hum MYL3 | 6xHis on MYL3 |
| Embryonic (Emb) | *MYH3* | Foetal skeletal and regenerating muscles |  | Endogenous Mouse | 6xHis |
| Perinatal (Peri) | *MYH8* | Foetal skeletal and regenerating muscles |  | Endogenous Mouse | 6xHis |
| Extraocular (Exoc) | *MYH13* | Extraocular and laryngeal muscles |  | None | e-GFP, 6xHis |
| Ila | *MYH2* | Fast skeletal | $1.401 \pm 0.101$ | Endogenous Mouse | e-GFP, 6xHis |
| Ilb | *MYH4* | Not expressed |  | Endogenous Mouse | e-GFP, 6xHis |
| Ild | *MYH1* | Fast skeletal | $3.022 \pm 0.935$ | Endogenous Mouse | e-GFP, 6xHis |

\* (18)
\*\*Endogenous mouse light chains are a combination of MLC1A, MLC3F, MLC1F, MLC2F
† (34)

Using MUSICO we recently re-examined the actin myosin-S1 ATPase cycle of fast rabbit muscle using both fast kinetic methods and steady-state ATPase assays to establish the primary parameters of an 8-step actin-myosin cross-bridge cycle (Fig 1; (9)). These parameters were then used with the addition of the overall ATPase parameters for the cycle, the k_cat_ and K_app_ (concentration of actin needed for half maximum ATPase rates) to model the complete cycle in solution. This allowed the occupancy of each state of the cycle to be predicted as a function of actin concentration. The occupancies calculated were then used to predict the duty ratio (DR; the proportion of the cycle myosin is bound to actin), the expected maximal velocity of contraction, (V_0_, in a motility assay or muscle sarcomere shortening) as a function of actin concentration and the effect of a 5 pN load on state-occupancies for a single motor. To test this approach we compared the information obtained for rabbit fast muscle myosin with two human cardiac myosin isoforms (α and β) that we had characterized previously (6). These illustrated how the cycle was altered to define myosins with different velocities and different sensitivities to load. While the duty ratio was largely unchanged among isoforms, the ATPase cycling rates and predicted velocities were altered in line with published experimental data. The effect of load reduced the predicted velocities and ATPase rates but to differing extents. Here, we have added in predictions about the economy of ATP usage during rapid shortening and whilst holding a 5 pN load.

**Figure 1.**
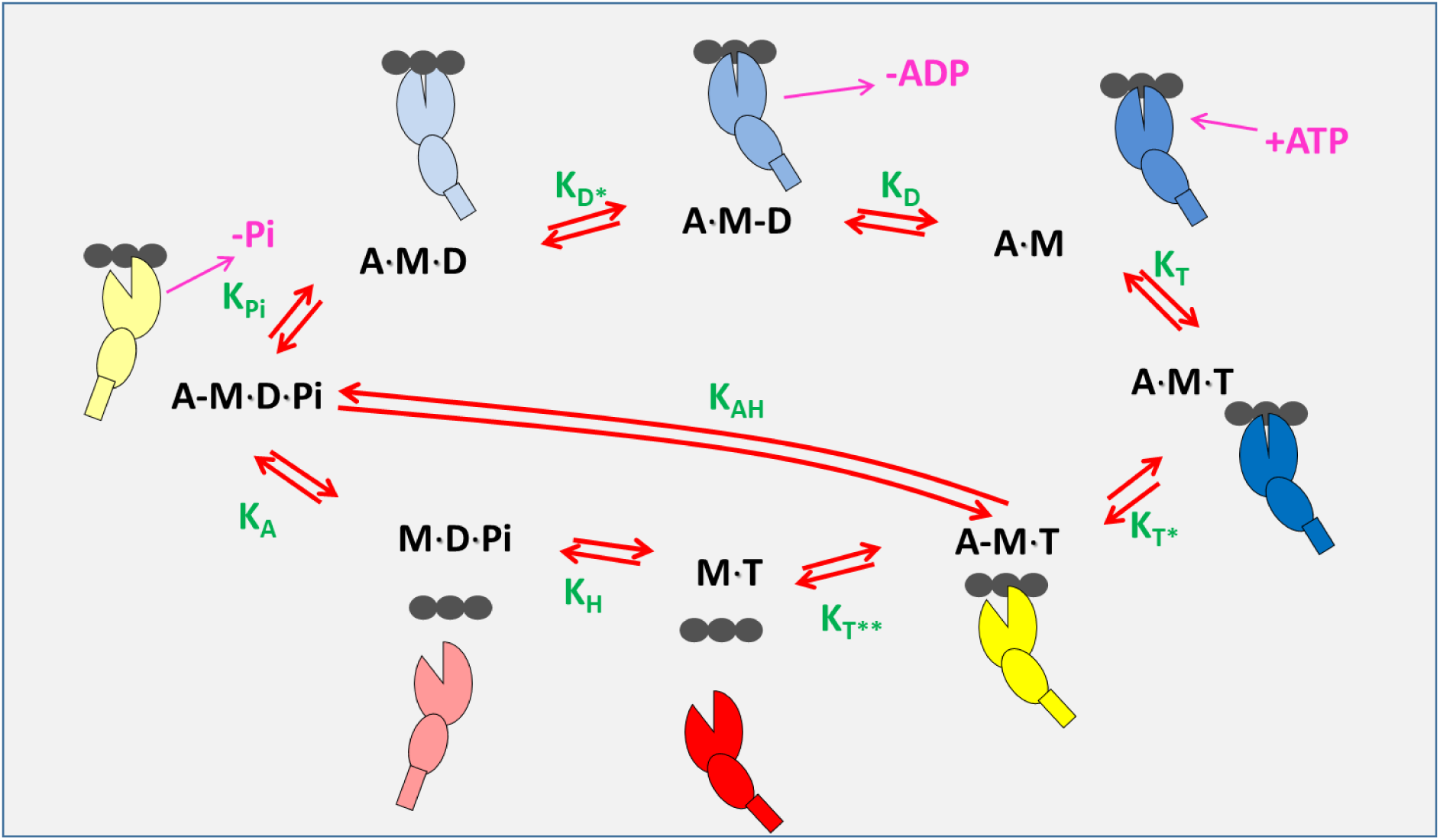
ATP driven actomyosin cross-bridge cycle. The black circles represent an actin filament composed of 3 actin monomers. Myosin is shown as 2 ellipses and a rod; the larger ellipse represents the upper and lower 50 k domains, the smaller ellipse and rod representing the converter domain, lever arm and light-chain binding region. Myosin in a strongly attached, force-holding state is represented in blue, weakly attached in yellow and detached in red shades. Green characters are the equilibrium constants for each step defined in the direction of ATP hydrolysis (clockwise).

In the current study, the analysis described above has been extended to four additional adult, fast-skeletal, human isoforms (IIa, IIb, IId and extraocular; ExOc) together with two developmental isoforms: embryonic (Emb) and perinatal (Peri) myosins. The analysis reveals that the relative velocities predicted for the isoforms vary widely (9-fold). Duty ratios vary over a narrow 2 fold range while economy of ATP usage varies 4-5 fold. Experimental data for these values are only available for a limited number of isoforms, but where available are compatible with our predictions. The extent to which these ATPase cycle studies can predict the properties of contracting muscle for both the well-defined and unstudied isoforms e.g. ExOc, Peri is discussed.

## Methods

### Protein expression and purification

Human muscle MyHC-sS1 for the β isoform and MyHC-S1 for the α isoform were expressed and purified as described previously (6). The motor domain of the β heavy chain was co-expressed with the N-terminal His_6_-tagged, human essential light-chain MYL3. The motor domain of the Emb myosin isoform was expressed with a His_6_-tag on the C-terminus (8). The fast muscle isoforms (IIb, IId, IIa), Peri, and ExOc isoforms were expressed with a C-terminally fused enhanced GFP and His_6_-tag. All proteins carrying a C-terminal His_6_-tag when purified carried the endogenous mouse light chains present in the C2C12 cells (see Table 1 (11). Briefly, replication incompetent recombinant adenoviruses were produced using the pAdEasy system containing expression cassettes encoding S1 of the human myosins under the transcriptional control of a CMV promoter. The adenoviral particles were amplified using HEK293 cells; the viruses were purified using CsCl gradients, and the concentrated virus was stored in a glycerol buffer at -20 °C. These adenoviruses were used to infect C_2_C_12_ myotubes in culture and cells were collected and frozen into cell pellets. Pellets were then homogenized in a low salt buffer and centrifuged, and the supernatants were purified by affinity chromatography using a HisTrap HP 1 ml column. The proteins were then dialyzed into the low salt experimental buffer (25 mM KCl, 20 mM MOPS, 5 mM MgCl_2_, 1mM DTT, pH 7.0).

Actin was prepared from rabbit muscle as described by (12). The actin was labelled with pyrene at Cys-374 as described in (13). When used at sub-micromolar concentrations the actin was stabilized by incubation in a 1:1 mixture with phalloidin.

### Kinetic measurements

Fast kinetic data for every isoform except for the Peri isoform have been previously published (6–8). All kinetic measurements for the Peri isoform were performed as previously described (6–8). Solutions were buffered with 20 mM MOPS, 5 mM MgCl_2_, 25 mM KCl, 1 mM DTT at pH 7.0, and measurements were conducted at 20 °C on a High-Tech Scientific SF-61 DX2 stopped-flow system. Traces were analyzed in Kinetic Studio (TgK Scientific) and Origin (OriginLab). The experimental data for all isoforms is summarized in Table S1. ATPase data for Peri, ExOc, IIa, IIb and IId isoforms were performed at 37 °C (11). To calculate the expected k_cat_ values at 20 °C, a Q_10_ value of 1.5 was used (14).

### Modeling

The published set of rate and equilibrium constants are summarized in Table S1, together with the k_cat_ and K_app_ values from steady-state actin-activated ATPase assays. With this data, the 8-state actin.myosin ATPase cycle was modelled using the MUSICO software (available at https://www.mijailovichlab.org/download) as described by (9, 10). The 8-step scheme of Fig 1 has a total of 24 rate and equilibrium constants but not all are independent. For each step, *i, K*_i_ = *k*_i_/*k*_*-i*_ and thus only two of the constants need to be defined experimentally for a complete description of the cycle. The free energy of ATP hydrolysis further constrains the overall balance of the cycle. Experiments have defined forward rate constants k_D*_, k_T*,_ k_H_, and the equilibrium constants K_T_and either K_D_ or K_D_K_D*_, in most cases to a precision of at least 20% (see Supplementary Table 1). The rate constants k_-T*_ and k_D_ are defined as diffusion limited. The events k_T**_ and k_-A_ are considered too fast to measure, and so have little effect on the modeling. Fitting the model to the actin-dependent steady-state ATPase data can give estimates for the equilibrium constants for actin binding (K_A_), ATP hydrolysis (K_H_) and on-actin hydrolysis step (K_AH_,) and the rate constants for phosphate release and ATP dissociation (k_Pi_ and k_T_ respectively). The K_i_ = k_i_/k_-i_ detailed balance equation can be used to define K_T*_, k_A_, k_-pi_, k_-D*_, k_-D_ and k_-T_, k_-H_ and k_-AH_. Initial concentration of ATP was set at 5 mM, and ADP and Pi at 0 mM; under steady-state conditions these are assumed to be zero.

The fraction of myosin in the strongly attached states AMD, AM-D, AM and AMT in the steady state, is defined as the *DR*. From *DR*, an estimate of the maximal velocity, *V*_o_, can be calculated from the equation *V*_o_=*d*/*τ*, where *d* is the distance over which myosin can produce force, and *τ* is the lifetime of the strongly attached state. The lifetime of the attached state is equal to *DR*/*ATPase rate*, hence, *V*_o_ = *d*·*ATPase*/*DR*. The economy of ATP usage at V_0_ at saturating actin concentration can be estimated V_0_ /ATPase from *DR* x *ATPase*/*d*.

In our previous modeling we used the data from two laboratories that used single molecule laser trap methods to define the effect of load on the ADP release step of the cycle (k_D*_; (15, 16)). These indicted that for β-cardiac a 5 pN load on actin.myosin slowed the ADP release by ∼3 fold. A similar effect on the power stroke (coupled to Pi release in our 8 state model) is also required to slow the ATPase cycling by ∼3-fold as reported for muscle fibers under isometric conditions. Here, in the absence of any direct measurements on any other isoform, we make the assumption that all isoforms have a similar load dependence. While this is an over simplification it does allows us to illustrate how load affects each isoform differently due to the changed balance of events in the cycle. To estimate the economy of ATP usage per pN of force generated at any actin concentration, the ATPase rate was divided by the load, here 5 pN.

The sensitivity matrices shown in Fig S3 demonstrate that with the exception of k_-T_, the fitted parameters are all well defined in the modeling program; values in the diagonal of >0.8 indicate well resolved parameters with little codependence. As reported previously (9), varying one of the fitted parameters (k_H_, k_-D_, K_D*_, k_-D*_, k_-T*_ or K_T*_) by ± 20 % has minimal effect on the best fit values for the remaining parameters. This observation remained true for the data presented in this study (see Table S4), with the remaining parameters varying by much less than 20%, with a few exceptions.

## Results & Discussion

We have modelled, using MUSICO, the complete ATPase cycle for all isoforms listed in Table 1. The modeling used the previously published values for the rate and equilibrium constants (Table S1) that have defined the cycle shown in Fig 1. The best fit parameters are listed in Table 2. In all cases, the measured constants were defined with a precision of at least ± 20%. The limitations of this precision on the modeling will be considered below, but in general, varying any of the parameters by ± 20% has a limited effect on the cycle; in most cases altering the occupancy of each state by much less than 20%. One new set of experimental data is presented here, that for the Peri isoform. Data collection was identical to that presented for all other isoforms, and the measured parameters are listed in Table S1.

**Table 2.**
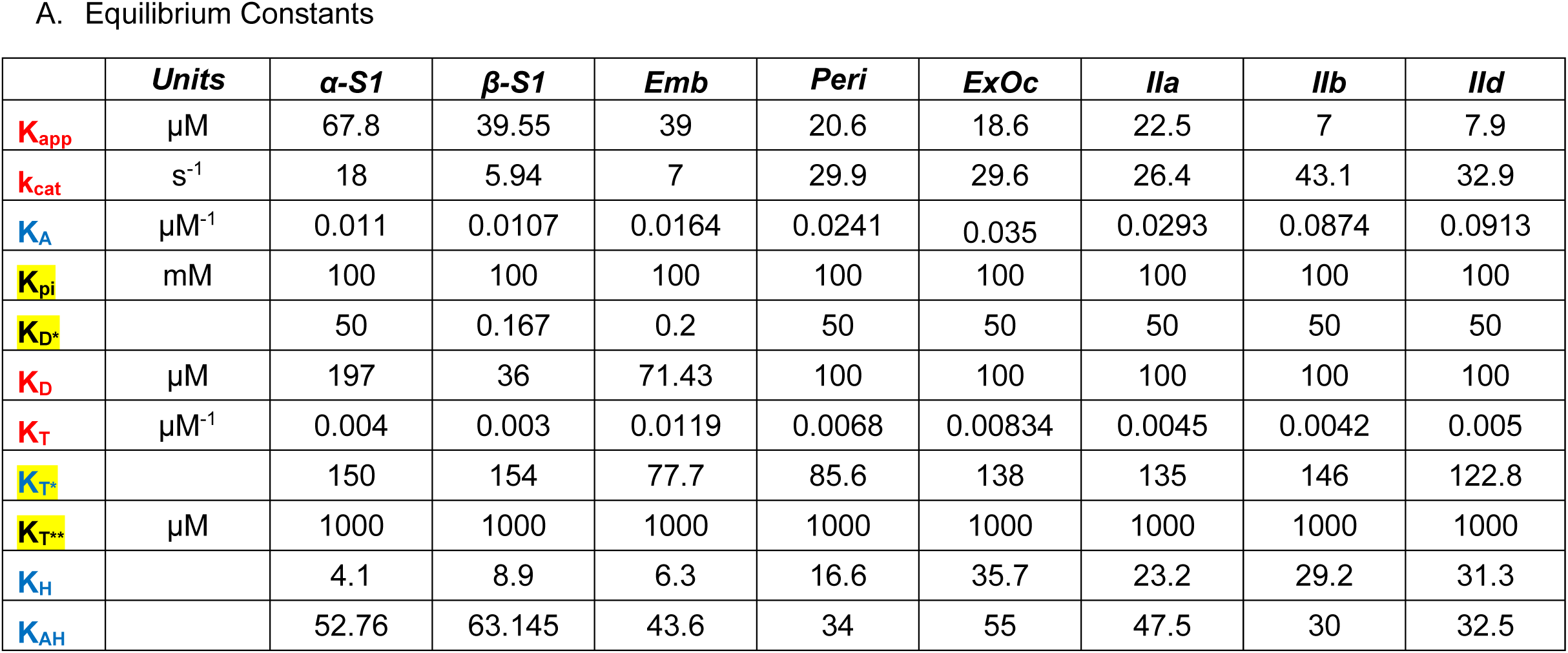
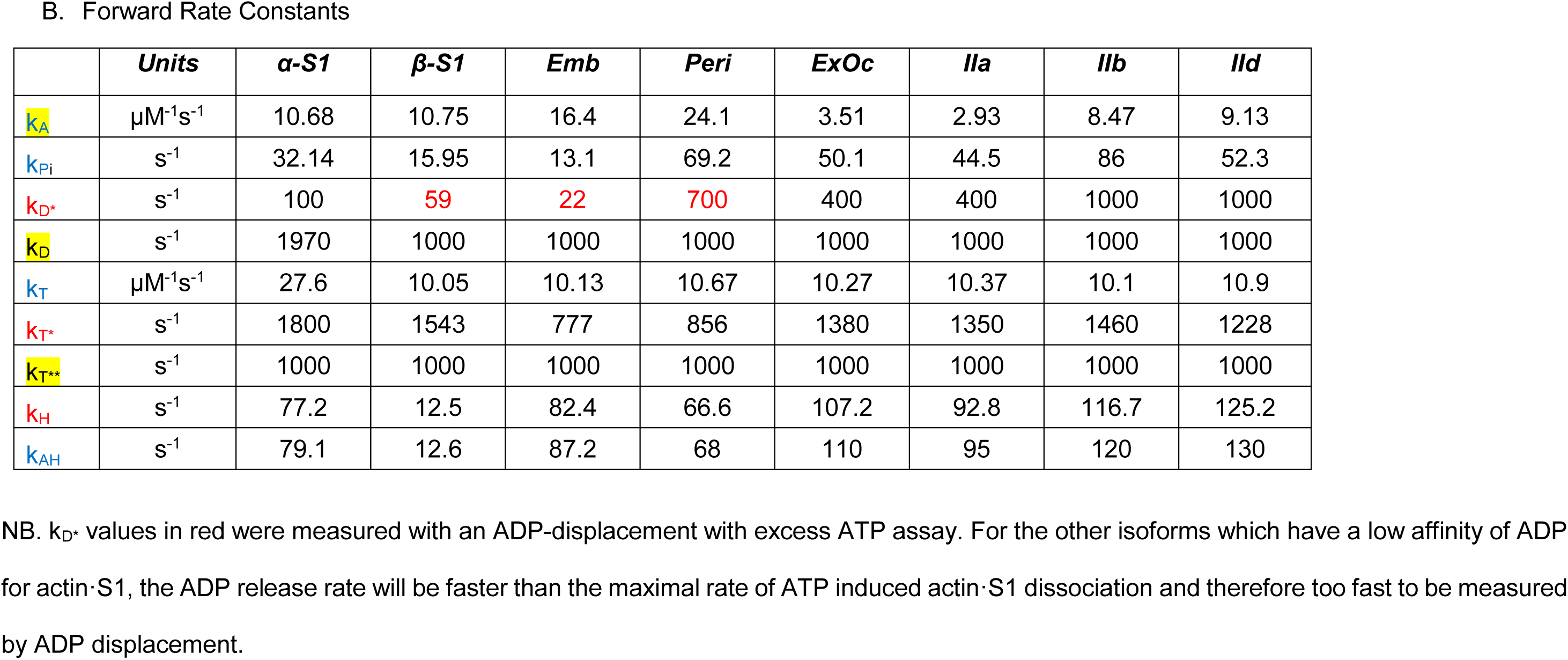
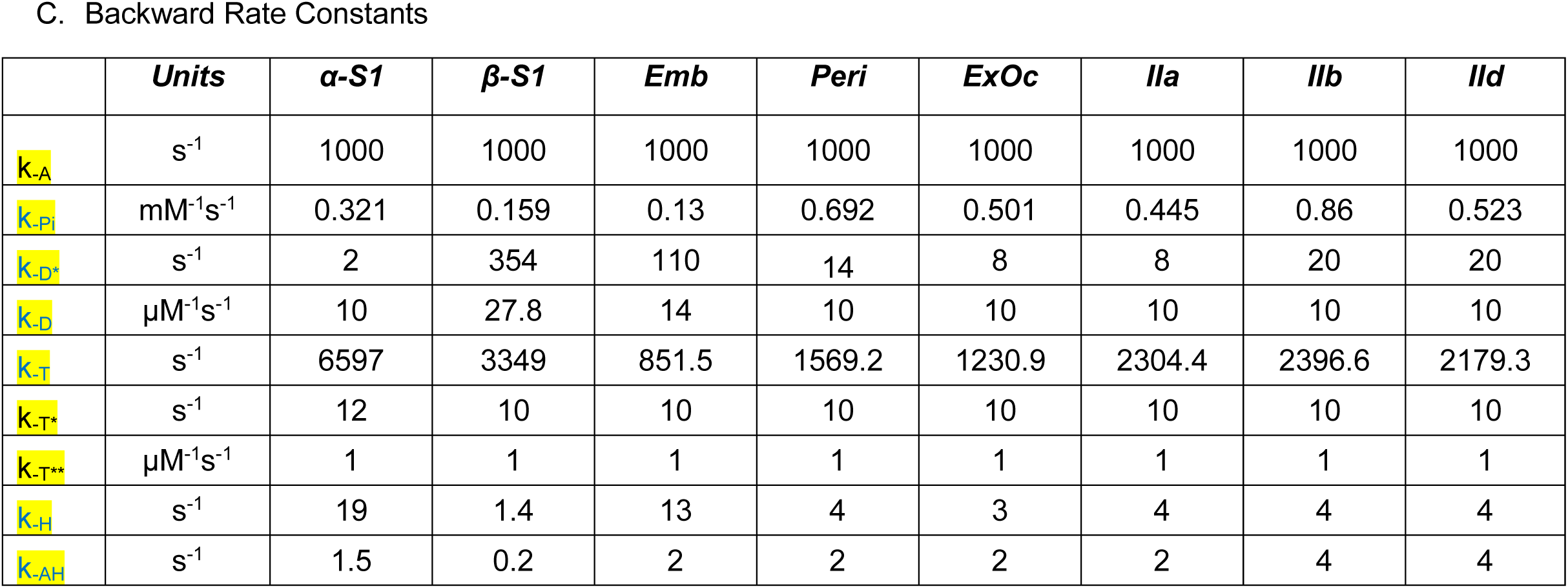
Fitted Rate and Equilibrium Constants of the ATPase cycle. Highlighted to show which are fitted, which are measured and which derived from assumption or detailed balance.

Using a combination of best fit and measured values for the cycle, the occupancy of each state in the cycle was calculated at three different actin concentrations [actin] = K_app_, (the actin concentration required for 50% of the V_max_ of the ATPase) and [actin] = 3·K_app_ and 20·K_app_ (actin concentration required for 75% and 95% of V_max_ respectively). A range of actin concentrations was chosen because it is not known what the appropriate actin concentration is in muscle fibers. Fig 2 presents the calculated occupancies for each state in the cycle as pie charts for each isoform. The color scheme of the pie charts matches that of the ATPase cycle states shown in Fig1, where red shades represent detached states, yellow shades weakly attached states and blue shades strongly attached states.

**Figure 2.**
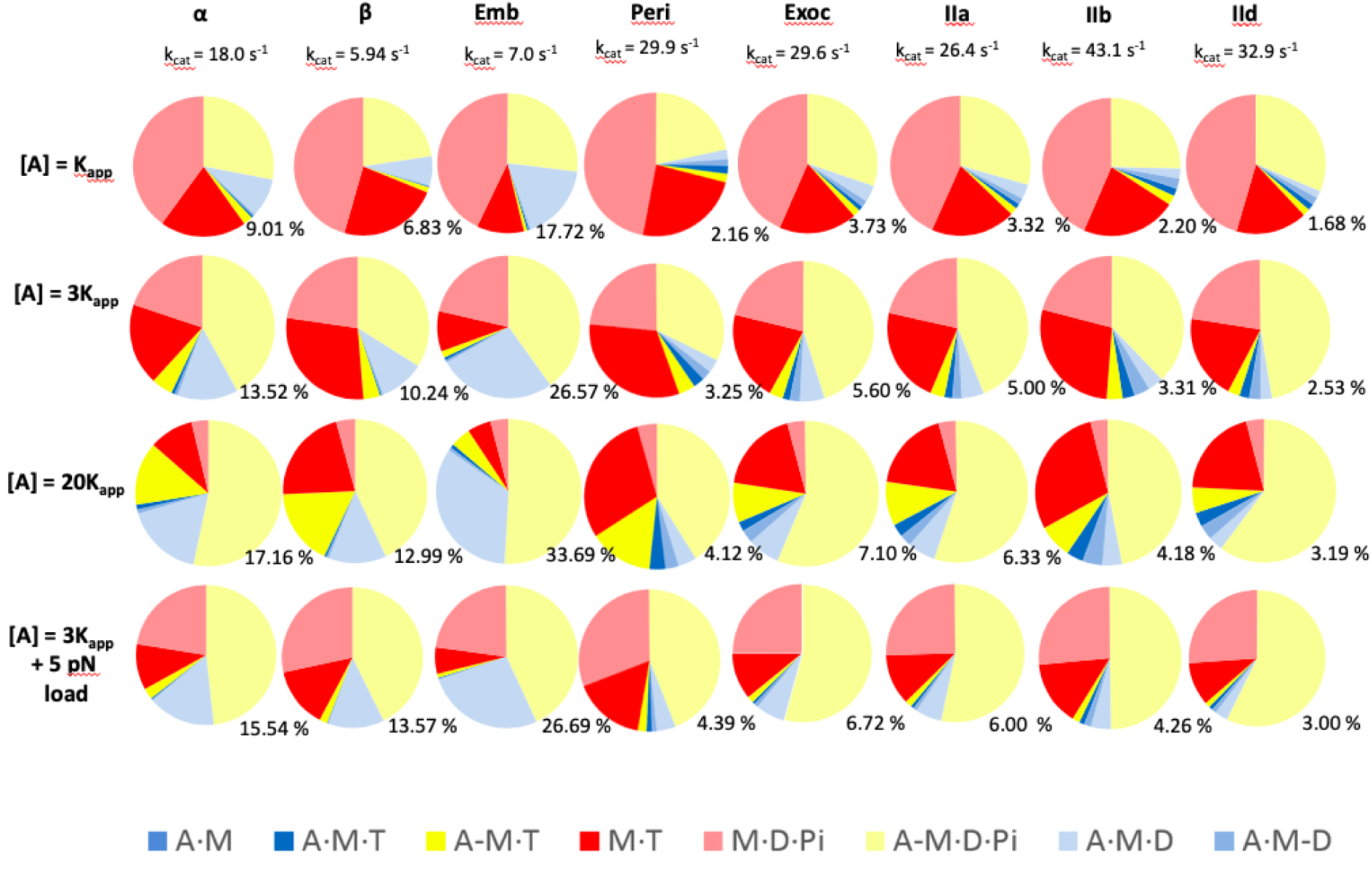
Fractional occupancies of each state in the ATPase at 3 different actin concentrations, [A] = *K*_m_, 3*K*_m_, and 20 *K*_m_ for each isoform and at [A] = 3Km plus 5 pN load. Colours of the Pie chart bans match those of Figure 1. The % value next to each chart gives the % of each cycle spent as the force holding A·M·D state.

### α & β Cardiac isoforms

For β-cardiac myosin, at low actin concentrations ([actin] = K_app_,), the detached state M·D·Pi, predominated (pale red shade; ∼45-50%), with similar amounts of the detached M·T and the weakly attached A-M·D·Pi state (each ∼25%). Only 6.8 % of the myosin is strongly attached as A·M·D, the predominant force holding state. All other species in the cycle have very low occupancy in the steady-state. As actin concentration increased (to 3 K_app_; 75 % V_max_ and 20K_app_; 95 % V_max_) the total occupancy of the detached states fell to ∼ 50% and then 25% and the weakly attached A-M·ATP and A-M·D·Pi increased from <25% to 35% and then >50% as expected. The strongly attached force holding states are dominated by the pale blue A·M·D state, which increases from 6.8% at low actin concentrations to 10% and 13.0 % as the actin concentration approached saturation. Thus, the DR is dominated by A·M·D and lies between 0.07 and 0.14, depending upon the actin concentration, at the zero load experienced in these solution assays. A similar pattern was observed for the human α-myosin except a slightly higher level of the A·M·D state, between 9.0% and 17.6% depending upon the actin concentration and a DR of 0.1 to 0-0.19.

Table 2 and Fig 2 list the measured k_cat_ values (ATPase cycling rates) for the α- and β-isoforms. Despite quite similar occupancies of the intermediates in the cycle, the cycling rates are very different; α-cardiac turns over ATP almost three times faster than β-cardiac. The predicted velocities were 2.25 fold faster for the α-vs the β-isoform (Fig 3, Table 3) which is similar to measured values (17). This calculation assumes an average 5 nm step for both. This result has implications for the economy of ATP usage which will be discussed below (see Fig 5).

**Table 3.**
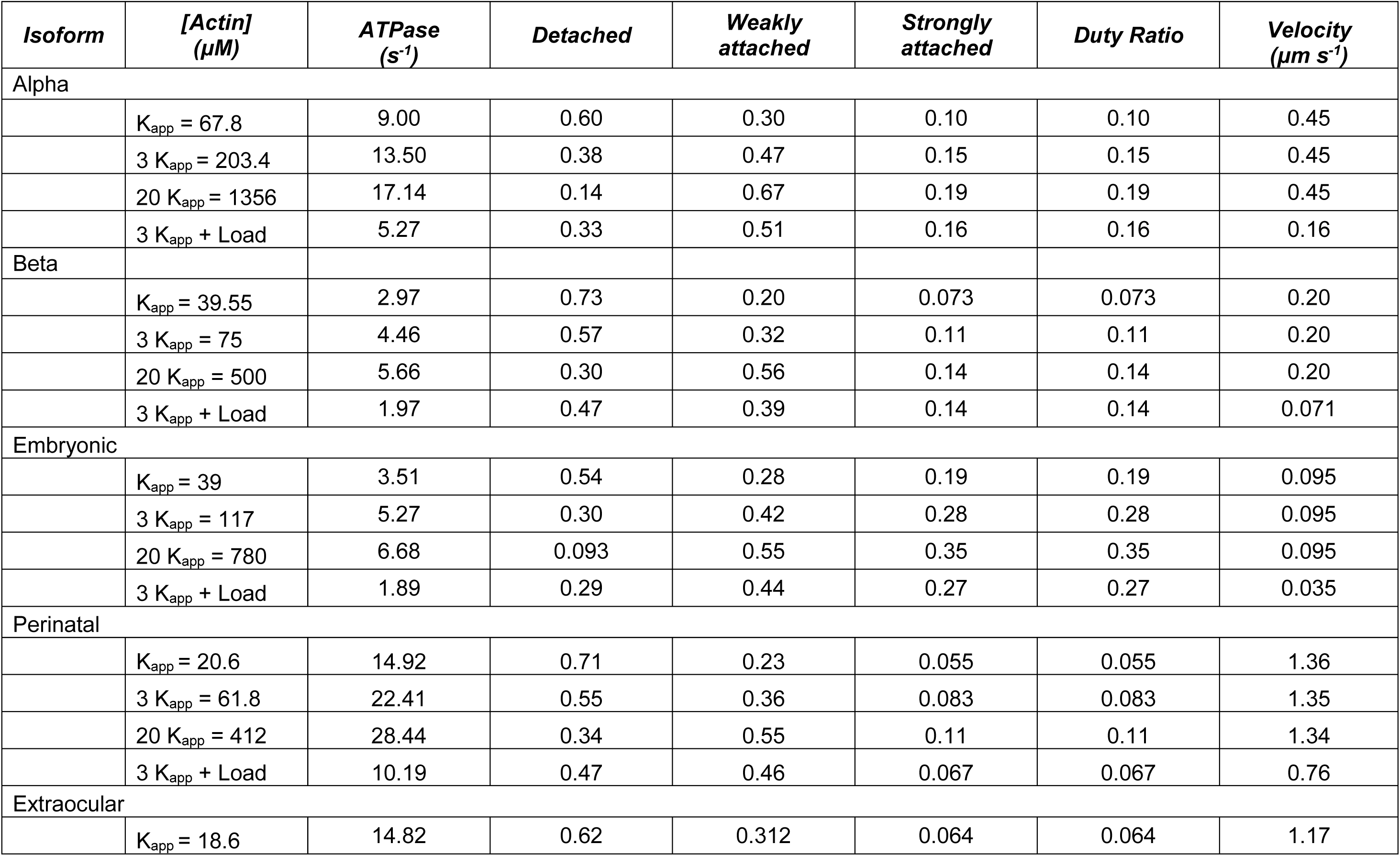
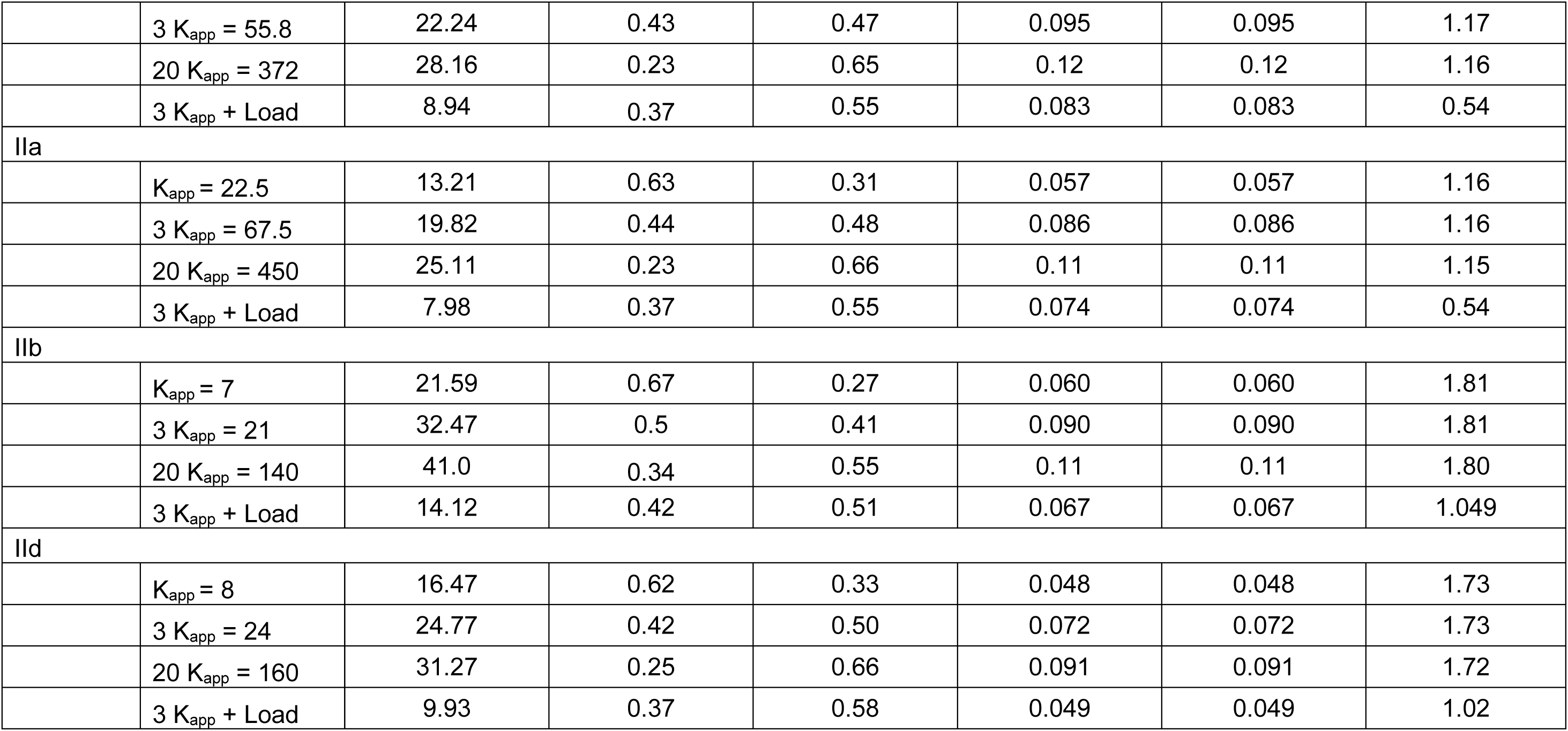
Predicted cross-bridge cycle parameters at three actin concentrations from the modelled ATPase cycle.

**Figure 3.**
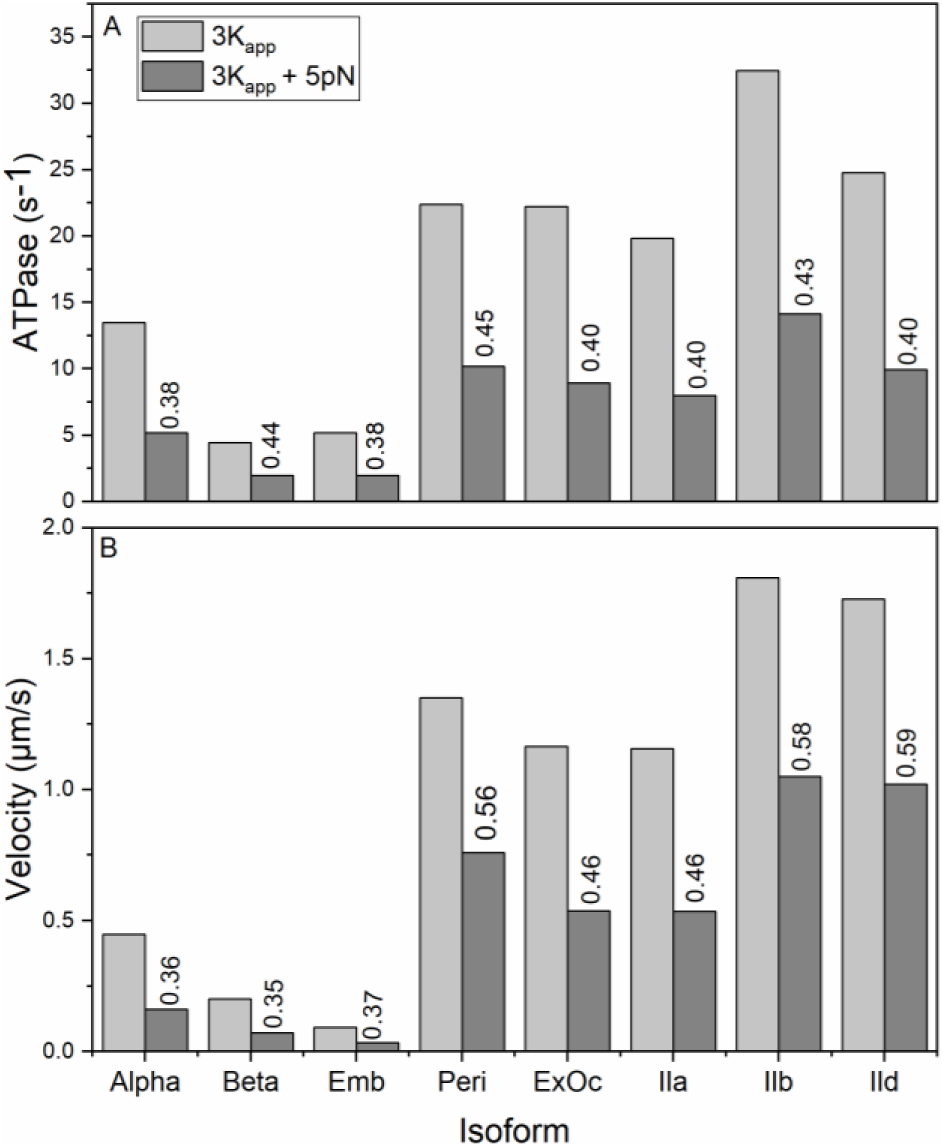
Plots of the estimated ATPase and V_0_ values at [actin] = 3 Km and the effect of a 5 pN load. The numbers give the loaded value as a fraction of the unloaded value.

**Figure 4.**
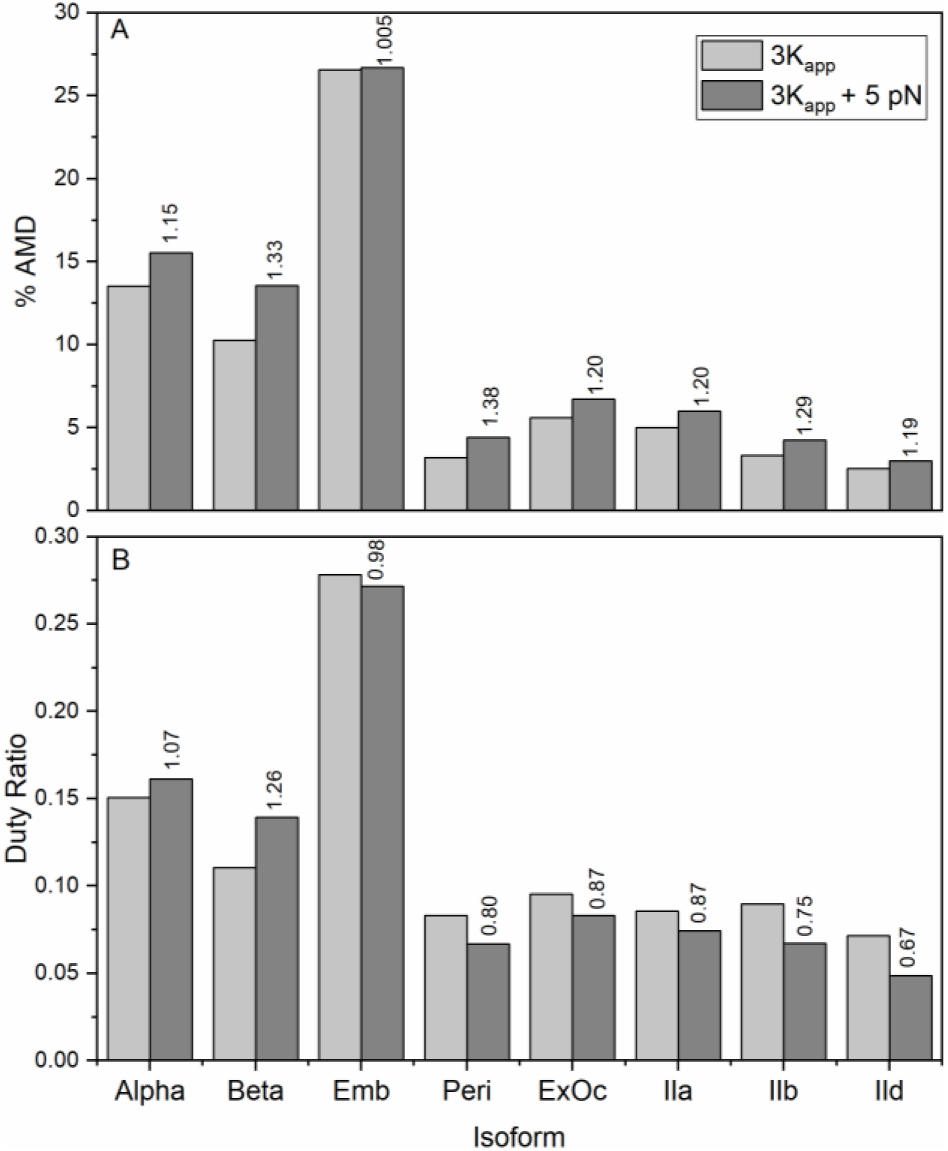
The estimated value of the Duty Ratio and the occupancy of the main force holding state A·M·D at [actin] = 3 Km and the effect of a 5 pN load. The numbers with each isoforms give the 5 pN loaded value as a fraction of the unloaded value.

**Figure 5.**
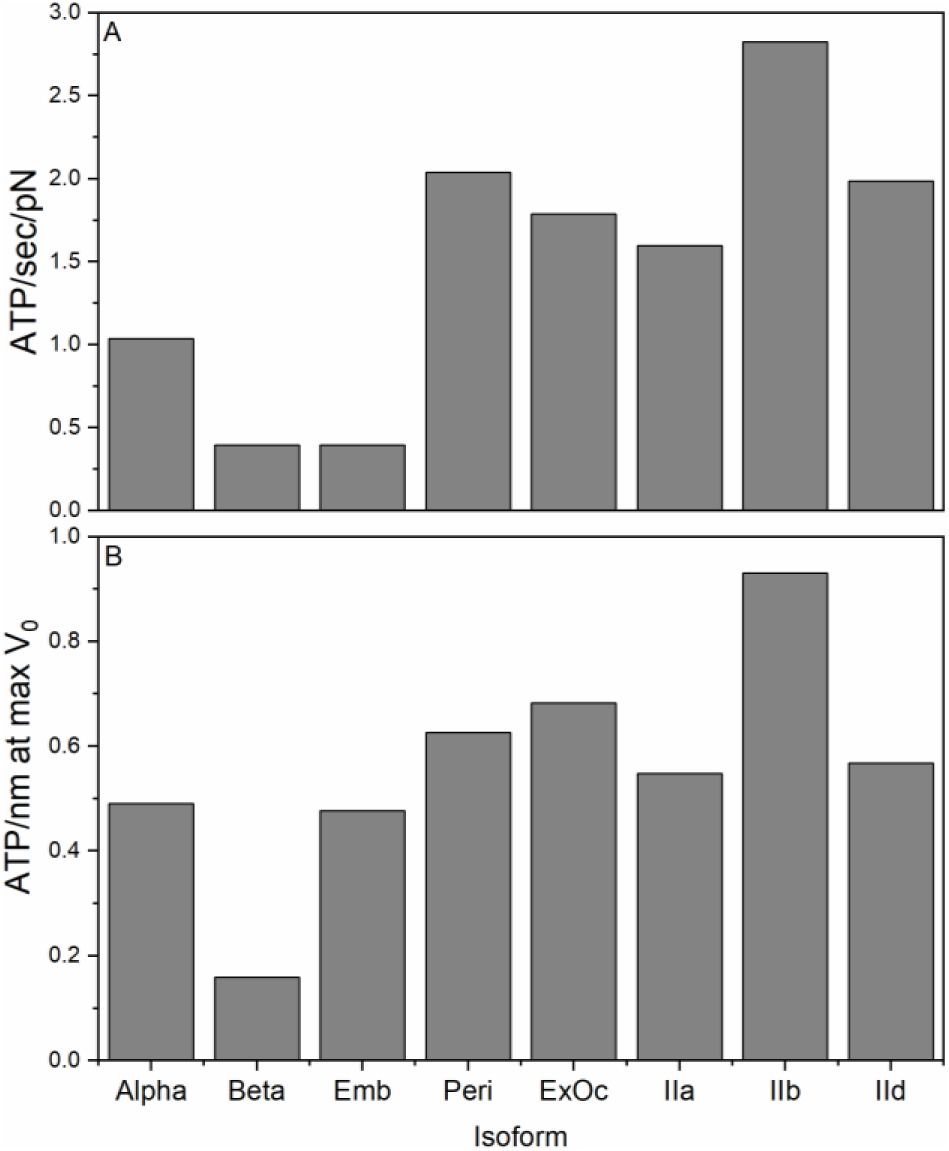
The economy of ATP usage (per myosin head), A) when holding a 5 pN load at [actin] = 3Km (assuming a similar load dependence for each isoform); B) when shortening at maximum velocity (V_0_) and V_max_ for the ATPase.

Single molecule laser trap assays have provided estimates of the load dependency of the ADP release rate constant (k_D*_) for human β myosin, and indicate that this rate constant slows down by ∼ a factor of 3 for a load of 5 pN (see methods (15, 16)). We argued that a similar load dependence is expected on the force generating step / Pi release step (k_Pi_ in Fig 1) and for the values used in the cycle a 3-fold reduction in k_Pi_ is required to slow the ATPase cycling rate under load, as has been reported for contracting muscle fibers (18, 19). Slowing both rate constants (k_Pi_ and k_D*_) down by a factor of 3 allows us to explore how the cycle would change under a 5 pN load, approaching the load which might be expected during isometric contraction of a fiber. The results are illustrated in the last row of the pie charts in Fig 2 for an actin concentration of 3K_app_ and the predicted effect on the ATPase rates and velocity of shortening are illustrated in Fig 3. There is no measurement of the load dependence of ADP release for any isoform other than β cardiac. For the purposes of illustration throughout we assume a similar load dependence for all isoforms to understand how a load may influence the cycle differently based on the observed differences in the cycle rate constants. The data for the effect of load on the state occupancy, the ATPase rate and the velocity is presented in Fig 2 & 3 and Table 3. For both α and β the ATPase cycling rates were reduced by approximately a factor of 3 while the occupancy of the A·M·D state increased from 10.2 to 13.6% for human β and 13.5 to 15 .5% for α. This implies that for an ensemble of myosins (in a thick filament or sarcomere) the α-myosin would maintain a higher occupancy of the force holding states. Note however, for β-myosin the occupancy of A·M·D increases by 1/3^rd^ under load, while for α-myosin, the occupancy increases by only ∼1/7^th^.

### Embryonic isoform

The complete experimental data set for the human Emb isoform was published in 2016 (8) and the published data are reproduced in Table S1. The results of the modeling are presented in Fig 2 - 4 in the same format as for the α- and β-isoforms for ease of comparison.

What is immediately striking in the Emb data set is that although the k_cat_ values for the Emb and β isoforms are similar (7.0 and 5.9 s^-1^ respectively) there are marked differences in the occupancy of the force holding A·M·D state (pale blue; Fig 1). While the detached M·D·Pi (pale red) and A-M·D·Pi (pale yellow) appear similar for β and Emb, the detached M·T (red) state is much smaller and the A·M·D state (pale blue) was much larger for Emb than for β (e.g. A·M·D = 26.5% for Emb vs 10.24% at [actin] = 3K_app_). This difference was even more marked when compared to the α-isoform. This large difference in A·M·D occupancy and duty ratio are brought about through differences in the contribution of three steps to the overall cycling speed (k_cat_): the hydrolysis step k_H_, the phosphate release step k_Pi_, and ADP release step k_D*_. For β myosin, k_H_ and k_Pi_ are comparable at 2-3 fold k_cat_ while k_D*_ is 10-fold larger than k_cat_ (see Table 4). Thus, as seen in Fig 2 for the β isoform at high actin concentration ([actin] = 20 K_app_), the predominant states are the ATP states M·T and A-M·T (together 45%), the weakly bound A-M·D·Pi (35%) and A·M·D (12.9%). In the case of Emb, k_cat_ is similar to that of β but the balance of the cycle is quite different: k_H_ is 10 k_cat_ and k_Pi_ and k_D*_ are now comparable at 2-3 fold k_cat_. So the ATP states M·T and A-M·T are much smaller while k_D*_, while the A-M·D·Pi (51%) and A·M·D (33%) states predominate, and thus a much larger DR is observed for Emb myosin.

**Table 4.** Balance of significant rate constants around the ATPase cycle.

| Isoform | $k_{cat}$ | $k_{Pi}/k_{cat}$ | $k_{D^*}/k_{cat}$ | $k_H/k_{cat}$ |
| --- | --- | --- | --- | --- |
| $\alpha$ | 18.00 | 1.78 | 5.56 | 4.28 |
| $\beta$ | 6.00 | 2.67 | 9.83 | 2.07 |
| Emb | 7.00 | 1.87 | 3.14 | 11.71 |
| Peri | 29.90 | 2.31 | 23.4 | 2.23 |
| Exoc | 29.60 | 1.69 | 13.51 | 3.72 |
| Ila | 26.40 | 1.74 | 15.15 | 3.60 |
| Ilb | 43.10 | 2.09 | 23.20 | 2.78 |
| Ild | 32.40 | 1.64 | 30.86 | 4.01 |

N.B. Slowest step (grey background) are in each case $\sim 2\times$ the $k_{cat}$ value. If this step is largely irreversible (true for $k_{Pi}$ not always true for $k_H$ ) then $\sim 50\%$ of the myosin will occupy the state before this slow step ( $k_{cat} = \theta \cdot k$ ; where $\theta$ is the fractional occupancy of that state and $k$ is the rate constant for the breakdown of that state).
2<sup>nd</sup> slowest step in all cases (yellow background) are $\sim 3\text{--}4$ times $k_{cat}$ and since $\theta = k_{cat}/k$ then $\theta \sim 0.25$ .
The third slowest step in each case is $\sim 10$ times $k_{cat}$ and so the occupancy of the relevant state will be approximately 0.1.

The higher occupancy of the force holding A·M·D state implies that the Emb isoform would be much better at holding loads than either of the two cardiac isoforms discussed so far. Assuming a similar load-holding capacity for each cross-bridge independent of the isoform, then there are almost twice as many cross-bridges present in the steady-state for Emb myosin, which suggests a fiber expressing Emb myosin would need to activate half the number of cross-bridges (per sarcomere or per thick filament) to hold the same load as a fiber expressing β myosin. At the whole fiber or whole muscle level, the differences in the packing of the filaments would need be considered.

In contrast to β or α, the presence of a 5 pN load on Emb has almost no effect on the occupancy of the force holding A·M·D state or the duty ratio. This was unexpected, but a closer examination of the effect of load suggests that since k_D*_ and k_Pi_ are both similar and dominate the ATPase cycling rate, when both are reduced to a similar extent by load, ATPase cycling is reduced by a factor of 3 but the balance of states around the cycle does not change significantly.

### Perinatal isoform

Like Emb, Peri is found in developing and regenerating muscle (2, 20). No biochemical kinetic study of this isoform has been published. Using the C2C12 expression system we have expressed the motor domain and completed a kinetic analysis as previously described for the other isoforms. Details of the measurements are given in Supplementary information (Fig S1-S4). The measured values for the steps in the ATPase cycle are listed in Table S1. The data do show distinct differences compared to the both the cardiac and Emb isoforms discussed so far.

As the Peri and Emb myosins are both developmental isoforms, a comparison between these is drawn here. A full comparison can be seen in Table S1. In the absence of actin the Peri S1 had an almost 3-fold slower second order rate constant of ATP binding to S1 compared to the Emb isoform (4.5 µM^-1^ s^-1^ vs 12.5 µM^-1^ s^-1^). The maximum rate of ATP binding (k_H_ + k_-H_) is almost 50% slower for the Peri than for the Emb (68.7 s^-1^ vs 130 s^-1^). This is assumed to measure of the ATP hydrolysis step. The Peri actin.S1 had a maximum rate of dissociation (k_+T*_) of 856 s^-1^, and is similar to the Emb S1 (777 s^-1^). However, the ATP binding affinity (K_T_), and hence the second order rate constant (K_T_k_+T*_), was almost two fold tighter than that of Emb (146.5 vs 84.3 µM Peri and Emb respectively). The crucial difference between the two developmental isoforms is the ADP release rate (k_+D*_) which is considerably faster (>700 s^-1^) than the Emb isoform (22 s^-1^). A slow ADP release rate is indicative of a slow type isoform such as β and Emb whereas the Peri is more like the fast skeletal isoforms (α, IIa, IIb, IId, ExOc). However, as shown in Fig S2 the rate constant for ADP release (k_D_ = 700 s^-1^) is only marginally slower than the maximum rate of actin dissociation by ATP (k_T*_ = 856 s^-1^). We have assigned this to k_D*_, but it could equally be assigned to k_D_.

The ATPase of the Peri isoform is much faster than Emb with a k_cat_ of almost 30 s^-1^ and the rapid release of ADP from A·M·D suggests a fast type myosin (21). The results of the modeling are shown in Fig 2-4.

Modeling the cycle shows a similar pattern to the α- and β-myosins but with some key differences. The occupancy of the force holding, pale blue, A·M·D state (pale blue) is much smaller for Peri (∼4.1% at [A] =20 K_app_) compared to α (17%) or β (13%) resulting in a smaller DR (0.11) than for β (0.14). The difference in DR could be considered small but examination of the pie charts in Fig 2 show a redistribution amongst the strongly attached states with the presence of significant amounts of the strongly attached states, A·M-D (∼2.8%) and A·M·T (∼3.5%) in addition to A·M·D. The presence of other strongly attached states appears to be a feature of the fast type myosins and will be discussed further when the fast adult myosins are considered.

Similar to the result for Emb isoform, the change in occupancy observed here for Peri is the result of changes in the balance between k_Pi_, k_D*_ and k_H_ and their contributions to k_cat_. While the rate of the hydrolysis step, controlled by k_H_, is much faster for Peri at 68.7 s^-1^ than for β (13.9 s^-1^), it is only twice the value of k_cat_. This means that the M·T and A-M·T states dominate at all actin concentrations considered, even more than is the case for β-myosin.

The estimate of velocity for this isoform is 1.35 µm.s^-1^ much faster than for α (0.45) or β (0.2). But this is dependent upon the assignment of k_D*_to 700 s^-1^. If instead this is k_D_ then there would be a missing value of k_D*_ which could be much lower than 700 s^-1^ and result in a much lower velocity.

### Fast skeletal isoforms

The adult fast skeletal isoforms, IIa, IIb and IId, form a closely related group of myosins and in humans they have ∼92 % sequence identity in the motor domain (22). The ExOc is also believed to be a fast muscle isoform based on contractile velocity of extraocular and pharyngeal muscles (23, 24). However, since it is only found in specialized muscle fibers expressing multiple isoforms, little is known about its biochemical and mechanical properties. It is ∼85 % identical to the adult fast isoforms. The experimental data for these four isoforms were published in two papers. Resnicow et al (11) published the first ever data on a set of recombinant human isoforms and Bloemink et al (7) followed this with a more detailed biochemical kinetic study of the same isoforms. The data in Table S1 is a summary of these studies.

Notably, like many larger mammals, humans have not been found to express any IIb protein although the gene is intact and theoretically capable of producing protein. The IIb we characterized is thus the only human IIb to have been studied. The fact that its properties appear similar to IIa and IId suggests that the gene has not degenerated significantly, despite not being expressed.

The k_cat_ values for the four isoforms (26 - 43 s^-1^) are similar to that of Peri (29.9 s^-1^) and much faster than the cardiac and Emb isoforms. As might have been expected, these four fast isoform show a very similar pattern of occupancy of the states in the cycle (Fig 2 & S2) with IIb being slightly different relative to the other three isoforms. The IIb isoform has a higher occupancy of the MT state and correspondingly lower A-M·D·Pi weakly attached state at high actin concentrations. All four have low occupancy of the A·M·D state (<10% in each case) but with significant variation amongst the 4, varying from 2.5% to 5.6% at [actin] = 3 K_app_. As seen for the Peri isoform the DR is not dominated by the A·M·D state; the other strongly attached states (A·M-D, A·M-T) contribute equally to the DR. We set the fast isomerization steps controlling ADP release and actin dissociation arbitrarily as a fast event at 1000 s^-1^. It is possible that, for these very fast myosin’s, this value of 1000 s^-1^ is too slow and should be considerably faster. We increased these values to 2000 or 3000 s^-1^ individually or as a group and repeated the modeling. This made little difference to the overall balance of the cycle (see Table S5 for the data set for myosin IId and IIb) with only those intermediates closely associated to the modified rate constant changed, if at all. For all other intermediates, the change was very small and always <10% of the initial value. The strongly attached states remained significantly occupied. The presence of these additional strongly attached states has implications for the DR and how sensitive the cycle is to load. This will be considered further below.

Before considering the implications of the modeling for understanding the different mechanochemical cycles and the role for which each isoforms has been optimized, we should first consider the limitations of the data sets used. There are limited amounts of the protein available for the assays used to define the ATPase cycle, and for good experimental reasons all data sets were not collected under identical conditions. The early experiments (all fast isoforms) were done at 20° C and 0.1 M KCl in order to make a better comparison with physiological conditions. Later, the conditions were changed to 25 mM KCl as this lower salt concentration was required to generate more reliable actin-activated ATPase data. The data presented for β myosin were collected at 20° C at both 100 and 25 mM KCl (6, 25) and allowed us to make corrections between salt conditions for each measured parameter. Similar studies have been published for the rabbit IIa isoform at both salt concentrations that supports the corrections made here (6, 9). The measured values are all listed in Table S1, and the corrected values are used in Table 2.

We state in the Methods section that we tested the robustness of our fitting by varying key fitted parameters by ± 20% and these have been published for the cardiac isoforms (9). The supplementary information (Table S4) shows representative data for the Emb isoform where one of K_D*_, k_D*_, K_T*_, k_T*_, k_-D_ or k_H_ were varied and all others refitted. Most parameters are change by very little; those that change by > 10% are highlighted in the Table S4 and are only those values directly linked to the altered parameter.

Having defined the ATPase & cross-bridge cycle for each of the 8 isoforms, we will now consider the implications of the different cycles for the contraction of muscle fibers containing each of these isoforms. Specifically, we will explore the maximum shortening velocity, the load-dependence of the cycle and the economy of ATP utilization.

### Maximum velocity of shortening, load dependence and economy of ATP usage

The maximum velocity of shortening (*V*_*o*_; zero load) of a muscle fiber expressing a single isoform can be estimated from the lifetime (τ) of the strongly attached force holding state (predominantly AMD) and the individual step size of the working stroke, *d*,

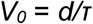

From our modeling τ can be calculated from *τ = DR/ATP*ase *rate* at any actin concentration. Since *DR* and *ATP*ase rates have very similar dependence on actin concentrations (both proportional to the fractional saturation of myosin with actin) *τ* and hence *V*_*0*_ are independent of actin concentration in this model. This means that the ATPase rate and velocity are not directly related except at saturating actin concentrations. The calculation of the *V*_*0*_ is identical for a contracting muscle fiber and velocity of actin movement in a motility assay, often measured in the absence of load. The model is therefore consistent with the observation that *V*_*0*_ is independent of the degree of activation of a muscle and only a very small number of myosin cross-bridges are required to achieve *V*_*0*_. For the purposes of the arguments set out below we have assumed the working stroke, *d*, to be 5 nm for each isoform.

The equation above assumes that the velocity is limited by the lifetime of the strongly attached states which, in the model used here, is controlled by the rate of cross-bridge detachment after completing the working stroke. This has been demonstrated to be true for the relatively slow β-type myosin where the ADP release rate constant (k_D*_) is easily measured. The same is true for the slow Emb isoform. For all other isoforms the ADP release rate constant has not been measured because either the rate constant is too fast for current methods or the relevant A·M·D complex cannot be easily formed by simply mixing ADP with A·M. The equilibrium K_D*_ in Fig 1 lies too far towards the A·M-D complex and little (<5%) A·M·D is formed. For fast rabbit muscle myosin the value of K_D*_was estimated as ∼50 (26). Thus we have good estimates of ADP release from β-cardiac and Emb myosin. For α, Peri and all fast muscle isoforms we have to estimate the ADP release based upon reasoned argument. If ADP release is too fast, then the lifetime and steady-state occupancy of the force holding state becomes too small and a muscle would be unable to hold much force. For example, a 1% occupancy of the force holding state would mean only 3 force holding cross-bridges for a fully activated 300 myosin thick filament. If the rate constant is too slow, the velocity becomes smaller than that observed experimentally. Here we have set that value for these fast fibers at the minimal possible, compatible with the expected velocities.

As noted above, fast muscle myosins have a higher predicted occupancy of other strongly attached states (dark blue); not just the A·M·D state. When k_D_ was doubled it had little effect on the occupancy of states in the cycle or the overall ATPase rates but velocity was increased by 15-20% and there was a 10% decline in the DR because of a ∼10% fall in the A·M-D state. The system does remain, however, a detachment limited model.

Fig 3B plots the predicted V_0_ values for each isoform and indicates there is an approximately 20 fold range of velocities from Emb (0.09 µm.s^-1^) to IId (1.66 µm.s^-1^). The order of predicted velocities and their values relative to Emb velocity is Emb [1], β [2], α [4], Peri [8], IIa and ExOc [12], IId and IIb [20]. Thus, our analysis of the cross-bridge cycle predicts Emb to be both the slowest of the isoforms and the one most capable of holding large steady state loads; i.e. longest lifetime of the A·M·D state and the highest occupancy of A·M·D in the steady state. In contrast, IId was the fastest isoform and has the lowest DR, and hence lowest force holding capacity.

The assumption of a 3 fold reduction in the rate constants for Pi and ADP release (k_Pi_ and k_D*_) induced by a 5 pN load predicts that for each isoform the different cycle characteristics result in different sensitivities of velocity to load. This effect of the cycle characteristics would remain true even if the measured load sensitivity varied for each isoform. The slower isoforms α, β and Emb have the highest sensitivity, with V_0_ being reduced by 2.7-2.8 fold. The velocity of ExOc, IIa and Peri are reduced 2.1 fold while the fastest isoforms IId and IIb show only a 1.6-1.7 fold reduction. This reflects the relative importance of the A·M-D and A·M-T strongly attached states. In the current model we have assumed that the fast events (rapid ADP release (K_D_), ATP binding (K_T_), and the ATP induced dissociation of actin (K_T*_ & K_T**_) are not directly affected by load.

The difference in load sensitivity of the cycle is reflected in the calculation of the economy of the isoforms shown in Fig 5 which plot ATP used per second when holding a force of 5 pN (Fig 5A) and when shortening at maximum velocity and saturating actin (zero load, Fig 5B). At a 5 pN load both β and Emb myosin have a similar economical usage of ATP (∼0.35 ATP/sec/pN) and are more economical than α (∼3 fold higher ATP usage) or the fast isoforms which use 1.5 to 2 fold more ATP than α myosin for the same load. In contrast, Emb is not as efficient in turnin*g* ATPase activity into movement as β myosin (0.2 ATP/sec per nm of travel) but is similar to alpha myosin which is uses just less than 0.5 ATP per nm. Thus Emb myosin appears to be designed for slow movement but economical force holding. As previously reported, this myosin is also able to continue functioning at much lower ATP concentrations than other myosin isoforms because of tight affinity for ATP (K_T_) (8). This property is shared with Peri and to a lesser extent with ExOc.

To fully understand the mechanical behavior of a muscle fiber, the force-velocity relationship is required as this can define the power output (Force x velocity) and the velocity at which the power output of the muscle is maximal. This is often considered to be the mechanical parameter that defines the optimal operating conditions for a muscle. An equivalent of the force-velocity curve at the single molecule level has recently been developed which shows how point mutations and small molecules can alter the relationship (27). Extrapolation between single molecule and whole fiber force-velocity curves is not trivial due to interactions between motors in the ensemble and the elasticity of the sarcomere filament. Thus our data cannot be used at present to predict the force-velocity relationship. However, the sarcomeric version of the MUSICO program is in principle capable of generating the force-velocity relationship (9). Modeling using this approach is a longer term project.

Our predictions can be compared, to a limited extent, to the published data on human muscle fibers where the myosin isoforms present are well defined. He et al (19) studied the force-velocity and ATPase properties of slow and type IIa human muscle fibers containing only β-myosin and myosin IIa respectively. Data were collected at 12 and 20 °C, and they calculated the economy of ATPase usage and the optimum velocities for power output. Pellegrino et al (18) additionally reported the velocities of human Type I, IIa and IId/x fibers at 12 °C (and velocities for the same set of fibers from mice, rat and rabbit). These studies therefore provide detailed muscle fiber data which can be compared to the predictions from our study. That said there are assumptions built in to any extrapolation from solution biochemistry to a muscle fiber that limits direct comparison. These include the number of myosin heads present and fully activated, in a muscle fiber, which in turn depends upon the density and packing of filaments in the muscle fiber. The units used to report velocities and ATPases differ due to these corrections. We will therefore limit ourselves to the values of the parameters relative to the values reported for Type 1 fibers containing β-myosin.

Table 5 lists the relative values of V_0_, ATPase and ATP economy under isometric conditions, provided by Pellegrino et al and He et al (18, 19). The relative values of V_0_ at 12 °C for Type IIa fibers relative to Type I were 4.2 for Pellegrino et al (18) and 2.4 for He et al. He et al had a similar ratio at 20 °C of 2.06 (19). Our data give a ratio nearer to that of Pellegrino of 5.8 for type IIa and 8.65 for type IId, compared to Pellegrino’s value of 9.15. Given the degree of error in each of these experimental measurements the values are of the correct order of magnitude. Similarly He et al measured the economy of ATP usage by Type I fibers to be 3.5 fold better than that of type IIa fibers. In contrast we calculate a 2.4 fold difference for the two types of myosin.

**Table 5.**
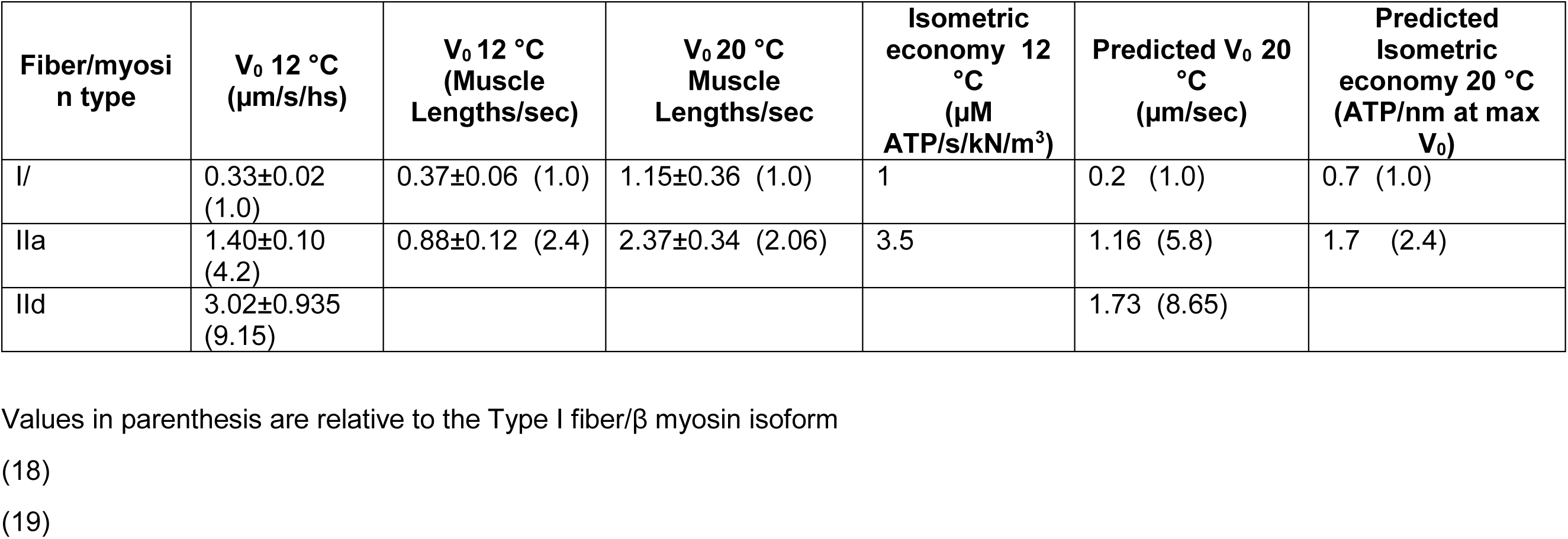
Comparison to Human muscle fiber data.

### Overview

Our modeling has revealed distinct characteristics of the ATPase cycle or each the human isoforms studied. Differences lie in the overall speed of the ATPase cycle (k_cat_) and the balance of the events in the cycle which effects how much time the myosin spends at each point of the cycle. There are three significant events in the cycle which define the characteristics of the cycle: 1) The Pi release step (k_Pi_) which controls entry into the strong actin-binding, force holding states. This event is shown as a single step but probably involves a myosin conformational change before or after the Pi release itself (28, 29). 2) The ADP release step controlled by the isomerization (k_D*_) which is followed by rapid ADP release, rapid ATP binding, and then actin dissociation. Thus, the ADP coupled isomerization is linked to detachment of the cross-bridge. 3) Finally, the ATP hydrolysis step which limits how long the cross-bridge remains detached before again being available to bind actin as A-M·D·Pi, which then gives access to the Pi release and force generating step. All other events are far more rapid and steps such as nucleotide binding/release and actin binding and release can be treated as rapid equilibration steps. Each of the myosins has a unique relationship between k_cat_ and the three events which is simply illustrated in Table 4 by showing the value of k_cat_ for each isoform and the value of each of the other rate constants relative to k_cat_. The values listed gives some an indication of the contribution of each transition to k_cat_. In all cases at least one of the three values is ∼ 2 times k_cat_, highlighted with a grey background in Table 4. This is k_Pi_ in most cases but for β and Peri the value of k_H_ is smaller or comparable to k_Pi_. A second value for each myosin is ∼3-5 times k_cat_ (yellow background). This is k_H_ for all fast isoforms and α, while it is k_D*_ for Emb and k_Pi_ for β and Peri. The third value is ∼ 10 times k_cat_ and this is k_D*_ in most cases except Emb. The value for α stands out as this is ∼5 times k_cat_ and similar to the value of k_H_. These different relationships between the three constants define the mechanical properties of the isoforms.

The isometric force of muscle fibers is normally found to be relatively invariant within the limits of the precision of the measurement and proportional to the number of strongly attached cross-bridges – although a contribution of weakly attached bridges cannot be ruled out. If this is true, then P_0_ will be a function of the number of active bridges and the DR. If all bridges are active (a function of Ca^2+^ activation, force activation, the super relaxed states and phosphorylation effects none of which operates in our pure S1 and actin system), then P_0_ is a function of DR. In our hands, the DR is a function of the actin concentration and the estimate of P_0_ will depend on the effective actin concentration present in the fiber. For a truly isometric fiber the actin concentration may not be the same for every myosin head due to the mismatch of the actin and myosin filament helices in a muscle. For this reason, we presented out data at different actin concentrations.

Our data show that the DR varies between isoforms: 0.05-0.075 for fast isoforms and 0.1-0.15 for slow/cardiac (even higher for Emb). Thus the expected P_0_ values per myosin head will be proportional to these DR values. Of course in the fiber the myosins act as an ensemble and the mechanical coupling between myosin may alter these numbers. In addition, while the density of thick filaments in a muscle fiber may be similar for all fast muscles, variations in packing are expected in slow and developing muscle where cell contents are not so exclusively packed with myofilaments.

The predicted V_0_ values vary in a way roughly compatible with expectation, the velocities are expected to be independent of actin concentration and so should be independent of the packing of filaments in the fiber. However, a question remains if unloaded shortening truly exists in the muscle fiber where some internal load may always be present, and therefore measured values will underestimate the true V_0_. Comparison of V_0_ muscle fiber values with motility velocities rarely show exact correspondence for reasons not yet fully explained. Although myosin orientation on the surface and the exact make up of actin filaments are thought to play a part in such discrepancies (30).

The solution data do not currently allow us to generate a force-velocity curve which would be required to define the optimal velocity for power output for each myosin type. This is believed to be the condition where the muscle is designed to operate. *Vo* and *Po* will provide the end points for the Force-Velocity curve but the shape of the relationship is distinct for different fibers and may depend on internal muscle elastic elements in addition to the ATPase cycle of different myosin motor domains. Studies of the load dependence of myosin isoforms using both single molecule methods and loaded motility assays should provide evidence that help define of the extent to which the information for the force velocity curve is contained within the myosin motor domains.

Our analysis of the differences in the cross-bridge cycle for each myosin isoform raises the issue of the sequence changes that bring about the adaptations to function. Earlier studies have examined groups of isoforms to identify key sequence changes (22, 31–33). These have often emphasised the variable surface loops in which isoform specific sequence changes occur. However, the source of the sequence changes required to bring about the changes in the overall balance of the cross-bridge cycle are likely to be more widespread. Future analyses will include a combination of bioinformatics, modeling and experimental investigation to define the sequences that generate the different properties of sarcomeric myosins and are responsible for their functional diversity.

## Acknowledgements

NIH GM29090 to LAL; NIH HL117138 to LAL. MAG & JW received funding from the European Union’s Horizon 2020 Research and Innovation Programme grant No. 777204, SILICOFCM.

## Author contributions

CAJ, MAG and LAL conceived the study. CAJ, with MS and SMM completed the kinetic modeling. JAW designed, performed and analyzed the stopped-flow experiments on the perinatal protein provided by CAV and AK. All authors contributed to the final version of the manuscript.

