## Supplementary files for "The ATPase cycle of Human Muscle Myosin II Isoforms: Adaptation of a single mechanochemical cycle for different physiological roles"

Running title: Human muscle myosin isoforms

*To whom correspondence should be addressed:

| | |
|---|---|
| Michael A Geeves | Leslie Leinwand |
| School of Biosciences, | BioFrontiers Institute and Department of Molecular, Cellular and Developmental Biology, |
| University of Kent, | University of Colorado Boulder, Boulder CO 80309 USA |
| Canterbury CT2 7NJ, UK | |
| m.a.geeves @kent.ac.uk | |
| +44 1227827597 | |

**Keywords:** developmental myosin, adult skeletal myosin, cardiac myosin, fast muscle, slow muscle, modeling, ATPase, ATP economy

### ***Supplementary Tables***

**Table S1. Experimentally measured constants for each isoform.**

| Step | Constant | *Alpha*[1] | *Beta*[1] | *Beta*[2] | *Embryonic*[3] | *Perinatal** | *Extraocula*[4] | *IIa*[4] | *IIb*[4] | *IId*[4] |
|---|---|---|---|---|---|---|---|---|---|---|
| | | *100 mM KCl* | *100 mM KCl* | *25 mM KCl* | *25 mM KCl* | *25 mM KCl* | *100 mM KCl* | *100 mM KCl* | *100 mM KCl* | *100 mM KCl* |
| ***ATP binding to S1*** | Second order rate constant of ATP binding ($\mu M^{-1}s^{-1}$) | 2.2 ± 0.1 | 1.5 ± 0.1 | 5.8 ± 0.4 | 12.5 ± 1.9 | 4.5 ± 1.0 | 1.0 ± 0.2 | 2.5 ± 0.5 | 1.7 ± 0.1 | 3.8 ± 0.2 |
| | $k_H + k_{-H}$ ($s^{-1}$) | 168 ± 28 | 14 | 91.2 ± 1.8 | 130 ± 3.4 | 68.7 ± 1.9 | 177 ± 8 | 141 ± 28 | 169 ± 2 | 195 ± 11 |
| ***ATP binding to actin.S1*** | $K_T k_{+T*}$ ($\mu M^{-1}s^{-1}$) | 2.5 ± 0.3 | 1.6 ± 0.3 | 4.4 ± 0.3 | 9.4 ± 1.0 | 5.9 ± 0.2 | 3.2 ± 0.4 | 1.7 ± 0.1 | 1.7 ± 0.3 | 1.7 ± 0.6 |
| | $K_T$ ($\mu M$) | 626 ± 143 | 710 ± 65 | 327.9 ± 53.3 | 84.3 ± 9.4 | 146.5 ± 9 | 360 | 662 | 715 | 602 |
| | $k_{+T*}$ ($s^{-1}$) | 1500 ± 167 | 1081 ± 50 | 1543 ± 100 | 777 ± 17 | 856 ± 26.9 | 1152 ± 60 | 1125 ± 78 | 1216 ± 283 | 1023 ± 215 |
| ***ADP affinity for actin.S1*** | $(K_{D*}+1)/K_D K_{D*}$ ($\mu M$) | 152 ± 25 | 10 ± 3 | 6.1 ± 0.7 | 14.3 ± 1.9 | 69.4 ± 11.2 | 352 ± 9 | 80 ± 15 | 54 ± 11 | 118 ± 33 |
| | $k_{+D*}$ ($s^{-1}$) | >1252 | 64 ± 3 | 58.7 ± 3.3 | 22.0 ± 1.8 | >700 | >1100 | >1100 | >1200 | >1000 |
| ***ATPase (0-10 mM KCl)*** | $k_{cat}$ ($s^{-1}$) | 18 | | 5.94 | 7.0 ± 0.105 | **59.7 ± 4.9 | **59.2 ± 5.7 | **52.8 ± 4.4 | **86.1 ± 8.4 | **65.7 ± 2.1 |
| | $K_{app}$ ($\mu M$) | 67.8 | | 39.55 | 38.5 ± 2.4 | 41.2 ± 3.7 | 37.2 ± 4.5 | 44.5 ± 5.6 | 14 ± 3.2 | 15.7 ± 1.3 |

[1] (6)
[2] (16)
[3] (8)
[4] (7)
* This study
** ATPase measurements conducted at 37 °C and 10 mM KCl (11)

**Table S2. Predicted Occupancy of the states during the ATPase cycle.**

| *Isoform* | *[Actin]* | *A·M* | *A·M·T* | *A-M·T* | *M·T* | *M·D·Pi* | *A-M·D·Pi* | *A·MD* | *A·M-D* |
|---|---|---|---|---|---|---|---|---|---|
| Alpha | | | | | | | | | |
| $K_{app}$ = 67.8 | $K_{app}$ | 0.0003 | 0.0051 | 0.021 | 0.20 | 0.40 | 0.28 | 0.090 | 0.0046 |
| | 3 $K_{app}$ | 0.0005 | 0.0078 | 0.048 | 0.18 | 0.20 | 0.42 | 0.14 | 0.0069 |
| | 20 $K_{app}$ | 0.0006 | 0.011 | 0.14 | 0.098 | 0.037 | 0.53 | 0.17 | 0.0087 |
| | 3 $K_{app}$ + Load | 0.0002 | 0.003 | 0.025 | 0.106 | 0.22 | 0.48 | 0.16 | 0.0026 |
| Beta | | | | | | | | | |
| $K_{app}$ = 39.55 | $K_{app}$ | 0.0002 | 0.002 | 0.014 | 0.28 | 0.45 | 0.19 | 0.068 | 0.003 |
| | 3 $K_{app}$ | 0.0032 | 0.0032 | 0.044 | 0.34 | 0.22 | 0.28 | 0.10 | 0.0045 |
| | 20 $K_{app}$ | 0.0004 | 0.005 | 0.21 | 0.26 | 0.042 | 0.36 | 0.13 | 0.0057 |
| | 3 $K_{app}$ + Load | 0.0001 | 0.0014 | 0.022 | 0.17 | 0.29 | 0.37 | 0.14 | 0.002 |
| Embryonic | | | | | | | | | |
| $K_{app}$ = 39 | $K_{app}$ | 0.00015 | 0.0046 | 0.0076 | 0.11 | 0.43 | 0.27 | 0.18 | 0.0035 |
| | 3 $K_{app}$ | 0.00022 | 0.0070 | 0.015 | 0.091 | 0.21 | 0.40 | 0.27 | 0.0053 |
| | 20 $K_{app}$ | 0.00029 | 0.0092 | 0.045 | 0.052 | 0.040 | 0.51 | 0.34 | 0.0067 |
| | 3 $K_{app}$ + Load | 0.00008 | 0.0025 | 0.0090 | 0.060 | 0.23 | 0.43 | 0.27 | 0.0019 |
| Perinatal | | | | | | | | | |
| $K_{app}$ = 20.6 | $K_{app}$ | 0.00080 | 0.018 | 0.019 | 0.24 | 0.47 | 0.22 | 0.022 | 0.015 |
| | 3 $K_{app}$ | 0.00120 | 0.027 | 0.040 | 0.32 | 0.23 | 0.32 | 0.032 | 0.022 |
| | 20 $K_{app}$ | 0.0016 | 0.035 | 0.14 | 0.30 | 0.044 | 0.41 | 0.041 | 0.028 |
| | 3 $K_{app}$ + Load | 0.00055 | 0.012 | 0.020 | 0.16 | 0.31 | 0.44 | 0.044 | 0.010 |
| Extraocular | | | | | | | | | |
| $K_{app}$ = 18.6 | $K_{app}$ | 0.00055 | 0.011 | 0.017 | 0.14 | 0.48 | 0.30 | 0.037 | 0.015 |
| | 3 $K_{app}$ | 0.00083 | 0.016 | 0.030 | 0.19 | 0.24 | 0.44 | 0.056 | 0.022 |

| | | | | | | | | | |
|---|---|---|---|---|---|---|---|---|---|
| | 20 $K_{app}$ | 0.0011 | 0.021 | 0.087961 | 0.18 | 0.045 | 0.56 | 0.071 | 0.028 |
| | 3 $K_{app}$ + Load | 0.000332 | 0.0066 | 0.013 | 0.087 | 0.28 | 0.54 | 0.067 | 0.0089 |
| IIa | | | | | | | | | |
| $K_{app}$ = 22.5 | $K_{app}$ | 0.0007 | 0.0099 | 0.016 | 0.15 | 0.48 | 0.30 | 0.033 | 0.013 |
| | 3 $K_{app}$ | 0.001 | 0.015 | 0.031 | 0.20 | 0.24 | 0.45 | 0.050 | 0.020 |
| | 20 $K_{app}$ | 0.0013 | 0.019 | 0.099 | 0.18 | 0.044 | 0.56 | 0.063 | 0.025 |
| | 3 $K_{app}$ + Load | 0.00042 | 0.0060 | 0.014 | 0.095 | 0.28 | 0.54 | 0.060 | 0.0080 |
| IIb | | | | | | | | | |
| $K_{app}$ = 7 | $K_{app}$ | 0.0011 | 0.015 | 0.021 | 0.19 | 0.48 | 0.25 | 0.022 | 0.022 |
| | 3 $K_{app}$ | 0.0017 | 0.022 | 0.035 | 0.26 | 0.23 | 0.38 | 0.033 | 0.032 |
| | 20 $K_{app}$ | 0.0022 | 0.029 | 0.075 | 0.29 | 0.043 | 0.48 | 0.042 | 0.041 |
| | 3 $K_{app}$ + Load | 0.00075 | 0.0098 | 0.017 | 0.13 | 0.29 | 0.49 | 0.043 | 0.014 |
| IId | | | | | | | | | |
| $K_{app}$ = 8 | $K_{app}$ | 0.00084 | 0.014 | 0.017 | 0.14 | 0.48 | 0.32 | 0.017 | 0.016 |
| | 3 $K_{app}$ | 0.0013 | 0.020 | 0.028 | 0.19 | 0.24 | 0.4 | 0.025 | 0.025 |
| | 20 $K_{app}$ | 0.0016 | 0.026 | 0.059 | 0.21 | 0.043 | 0.60 | 0.032 | 0.031 |
| | 3 $K_{app}$ + Load | 0.00051 | 0.0082 | 0.013 | 0.093 | 0.28 | 0.57 | 0.030 | 0.0099 |

**Table S3. Resolution matrices for the embryonic and perinatal isoforms.**

| **Embryonic** | **Predicted value** | $\mathbf{K_A}$ | $\mathbf{k_{pi}}$ | $\mathbf{k_{-T}}$ | $\mathbf{K_H}$ |
|---|---|---|---|---|---|
| $\mathbf{K_A}$ | 0.0164 | **0.99** | 0.0047 | -0.00024 | -0.0021 |
| $\mathbf{k_{pi}}$ | 13.1 ($s^{-1}$) | 0.0047 | **0.98** | 0.00025 | 0.081 |
| $\mathbf{k_{-T}}$ | 851.5 ($s^{-1}$) | 0.00024 | 0.00025 | **0.00000032** | -0.00001 |
| $\mathbf{K_H}$ | 6.34 | 0.0021 | 0.082 | -0.000011 | **0.88** |
| $\mathbf{K_{AH}}$ | 43.6 | 0.034 | 0.090 | 0.000067 | -0.45 |

| **Perinatal** | **Predicted value** | $\mathbf{K_A}$ | $\mathbf{k_{pi}}$ | $\mathbf{k_{-T}}$ | $\mathbf{K_H}$ |
|---|---|---|---|---|---|
| $\mathbf{K_A}$ | 0.019 | **0.99** | 0.000030 | -0.0013 | -0.000014 |
| $\mathbf{k_{pi}}$ | 89.9 ($s^{-1}$) | 0.000030 | **0.99** | 0.0014 | 0.00011 |
| $\mathbf{k_{-T}}$ | 584.7 ($s^{-1}$) | -0.0013 | 0.0014 | **0.0000036** | -0.000004 |
| $\mathbf{K_H}$ | 15.6 | -0.000014 | 0.00011 | -0.0000037 | **0.99** |

**Table S4. Sensitivity analysis; Percentage change of the predicted constants induced by a change of +20% or -20% to one of the six fitted parameters for the embryonic isoform.**

Red numbers indicate the % change imposed in each set, Blue/yellow background colors indicate modelled constants that decrease (blue) or increase (yellow) by > 10% (pale shade) or 20% (dark).

| | ***Units*** | ***Estimate*** | ***$K_{D^*}$+20%*** | ***$K_{D^*}$-20%*** | ***$k_{D^*}$ +20%*** | ***$k_{D^*}$ -20%*** | ***$K_{T^*}$ +20%*** | ***$K_{T^*}$-20%*** | ***$k_{T^*}$ +20%*** | ***$k_{T^*}$ -20%*** | ***$k_{-D}$ +20%*** | ***$k_{-D}$ -20%*** | ***$k_H$ +20%*** | ***$k_H$ -20%*** |
|---|---|---|---|---|---|---|---|---|---|---|---|---|---|---|
| ***Equilibrium Rate Constants*** | | | | | | | | | | | | | | |
| $K_A$ | $\mu M^{-1}$ | 0.02 | 10.37 | -16.46 | 9.76 | -14.63 | 0.00 | -0.61 | 0.00 | -0.61 | 1.22 | -1.83 | 0.61 | -1.22 |
| $K_{D^*}$ | | 0.20 | 20.00 | -20.00 | 0.00 | 0.00 | 0.00 | 0.00 | 0.00 | 0.00 | 0.00 | 0.00 | 0.00 | 0.00 |
| $K_D$ | (μM) | 71.43 | 0.00 | 0.00 | 0.00 | 0.00 | 0.00 | 0.00 | 0.00 | 0.00 | -16.67 | 25.00 | 0.00 | 0.00 |
| $K_{T^*}$ | | 77.70 | 0.00 | 0.00 | 0.00 | 0.00 | 20.00 | -20.00 | 0.00 | 0.00 | 0.00 | 0.00 | 0.00 | 0.00 |
| $K_H$ | | 6.34 | 3.38 | -5.38 | 4.50 | -1.38 | 0.99 | -0.41 | 0.14 | -0.22 | -0.45 | -0.98 | 20.00 | -20.00 |
| $K_{AH}$ | | 43.62 | 3.43 | -5.44 | 4.62 | -1.26 | 1.10 | -0.10 | 0.14 | -0.22 | -0.48 | -1.00 | 21.23 | -20.82 |
| ***Forward Rate Constant*** | | | | | | | | | | | | | | |
| $k_A$ | $\mu M^{-1}s^{-1}$ | 16.40 | 10.37 | -16.46 | 9.76 | -14.63 | 0.00 | -0.61 | 0.00 | -0.61 | 1.22 | -1.83 | 0.61 | -1.22 |
| $k_{Pi}$ | $s^{-1}$ | 13.10 | -10.69 | 19.85 | -9.92 | 16.79 | -0.38 | 0.00 | -0.76 | 0.00 | -1.53 | 1.53 | -3.05 | 4.58 |
| $k_{D^*}$ | $s^{-1}$ | 22.00 | 20.00 | -20.00 | 20.00 | -20.00 | 0.00 | 0.00 | 0.00 | 0.00 | 0.00 | 0.00 | 0.00 | 0.00 |
| $k_T$ | $\mu M^{-1}s^{-1}$ | 10.13 | -0.04 | -0.01 | -0.02 | -0.08 | 0.02 | -0.01 | 0.00 | 0.00 | -0.01 | -0.01 | -0.10 | -0.04 |
| $k_{T^*}$ | $s^{-1}$ | 777.00 | 0.00 | 0.00 | 0.00 | 0.00 | 20.00 | -20.00 | 20.00 | -20.00 | 0.00 | 0.00 | 0.00 | 0.00 |
| $k_H$ | $s^{-1}$ | 82.38 | 3.38 | -5.38 | 4.50 | -1.38 | 0.99 | -0.41 | 0.14 | -0.22 | -0.45 | -0.98 | 20.00 | -20.00 |
| $k_{AH}$ | $s^{-1}$ | 87.24 | 3.43 | -5.44 | 4.62 | -1.26 | 1.10 | -0.10 | 0.14 | -0.22 | -0.48 | -1.00 | 21.23 | -20.82 |
| ***Backward Rate Constants*** | | | | | | | | | | | | | | |
| $k_{-Pi}$ | $mM^{-1}s^{-1}$ | 0.159 | -10.69 | 19.85 | -9.92 | 16.79 | -0.38 | 0.00 | -0.76 | 0.00 | -1.53 | 1.53 | -3.05 | 4.58 |
| $k_{-D}$ | $\mu M^{-1}s^{-1}$ | 14.00 | 0.00 | 0.00 | 0.00 | 0.00 | 0.00 | 0.00 | 0.00 | 0.00 | 20.00 | -20.00 | 0.00 | 0.00 |
| $k_{-T}$ | $s^{-1}$ | 851.50 | -0.04 | -0.01 | -0.02 | -0.08 | 0.02 | -0.01 | 0.00 | 0.00 | -0.01 | -0.01 | -0.10 | -0.04 |
| $k_{-T^*}$ | $s^{-1}$ | 10.00 | 0.00 | 0.00 | 0.00 | 0.00 | 0.00 | 0.00 | 20.00 | -20.00 | 0.00 | 0.00 | 0.00 | 0.00 |

**Table S5. The effect of increasing the fast isomerisation forward and backward rate constants by a factor of 2 on the occupancies of states in the cycle.**

| [Actin] | Isoform | Step changed | M·DPi | A-M·D·Pi | A·M·D | A·M-D | A·M | A·M·T | A-M·T | A-M·T | ATPase ($s^{-1}$) | Velocity (*$\mu m\ s^{-1}$*) | Duty Ratio |
|---|---|---|---|---|---|---|---|---|---|---|---|---|---|
| $3K_{app}$ | IIb | None | 0.23 | 0.38 | 0.033 | 0.032 | 0.0017 | 0.022 | 0.035 | 0.26 | 32.47 | 1.808 | 0.090 |
| | | $2k_D$ | 0.238 | 0.384 | 0.0033 | 0.0165 | 0.00174 | 0.0229 | 0.0357 | 0.267 | 33.01 | 2.217 | 0.074 |
| | | $2k_{T^{**}}$ | 0.235 | 0.377 | 0.0033 | 0.0324 | 0.00171 | 0.0224 | 0.0214 | 0.277 | 32.43 | 1.81 | 0.090 |
| | | 2x both | 0.239 | 0.383 | 0.033 | 0.0164 | 0.00174 | 0.0227 | 0.0217 | 0.282 | 32.98 | 2.22 | 0.074 |
| $3K_{app}$ + 5 pN load | IIb | None | 0.29 | 0.49 | 0.043 | 0.014 | 0.00075 | 0.0098 | 0.017 | 0.13 | 14.12 | 1.049 | 0.067 |
| | | $2k_D$ | 0.295 | 0.496 | 0.00428 | 0.00711 | 0.00033 | 0.00986 | 0.0169 | 0.132 | 14.23 | 1.183 | 0.060 |
| | | $2k_{T^{**}}$ | 0.293 | 0.492 | 0.00426 | 0.0141 | 0.00033 | 0.00974 | 0.0102 | 0.137 | 14.11 | 1.056 | 0.067 |
| | | 2x both | 0.295 | 0.496 | 0.0428 | 0.00711 | 0.0033 | 0.00981 | 0.0103 | 0.138 | 14.22 | 1.184 | 0.060 |
| $3K_{app}$ | IId | None | 0.24 | 0.474 | 0.025 | 0.025 | 0.0013 | 0.020 | 0.028 | 0.19 | 24.77 | 1.728 | 0.072 |
| | | $2k_D$ | 0.238 | 0.480 | 0.0253 | 0.0125 | 0.00129 | 0.0207 | 0.0279 | 0.194 | 25.09 | 2.10 | 0.060 |
| | | $2k_{T^{**}}$ | 0.236 | 0.473 | 0.0252 | 0.0248 | 0.00127 | 0.0203 | 0.0169 | 0.203 | 24.75 | 1.729 | 0.072 |
| | | 2x both | 0.239 | 0.479 | 0.0253 | 0.0125 | 0.00128 | 0.0206 | 0.0171 | 0.205 | 25.07 | 2.10 | 0.060 |
| $3K_{app}$ + 5 pN load | IId | None | 0.28 | 0.57 | 0.030 | 0.0099 | 0.00051 | 0.0082 | 0.013 | 0.093 | 9.93 | 1.021 | 0.049 |
| | | $2k_D$ | 0.277 | 0.573 | 0.0300 | 0.0050 | 0.00051 | 0.00823 | 0.0128 | 0.0936 | 9.98 | 1.140 | 0.044 |
| | | $2k_{T^{**}}$ | 0.276 | 0.570 | 0.0300 | 0.0099 | 0.00051 | 0.00815 | 0.00786 | 0.0982 | 9.93 | 1.022 | 0.049 |
| | | 2x both | 0.277 | 0.573 | 0.0300 | 0.0050 | 0.00051 | 0.00819 | 0.00790 | 0.0987 | 9.98 | 1.141 | 0.044 |

Grey background highlights any value that changes by > 10%. Note these are all very low occupancy states that have little consequence for the overall ATPase rates or on the occupancy of most other states. Increases in $k_D$ do cause a ~15-20% increase in $V_0$ and a ~10% decrease in DR.

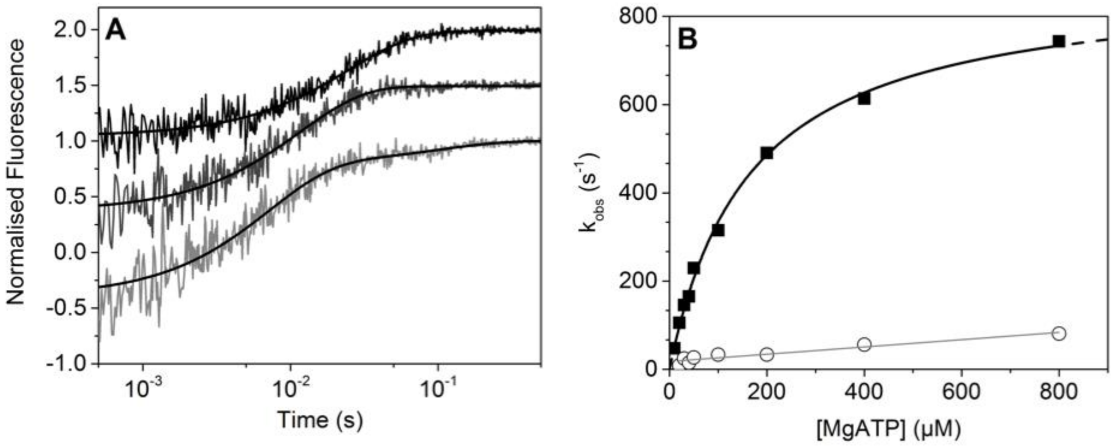

**Figure S1.** ATP induced dissociation of Perinatal S1 from pyrene-labelled actin in the absence and presence of ADP. (A) Example transients of ATP induced dissociation of S1 from pyrene-labelled actin. Transients have been off-set on the y-axis for clarity. The concentration of S1.pyrene-labelled actin complex was kept constant at 50 nM while the ATP concentration was increased. The example transients here show 10, 20 and 30 μM ATP. At ATP concentrations above 10 μM the transients were best described by a double exponential. The $k_{obs}$ of both phases of the transient were plotted in B. (B) When the $k_{obs}$ is plotted against the ATP concentration the fast phase had a hyperbolic dependence giving a maximum $k_{obs}$ = $k_{+T*}$ of 883.9 ± 21.9 $s^{-1}$, the concentration for half-maximum $k_{obs}$ = $K_T$ of 163.9 ± 10.5 μM, and the second-order rate constant $K_T k_{T*}$ of 5.4 ± 0.24 μM. These were repeated three times and the average is summarised in Table S1.

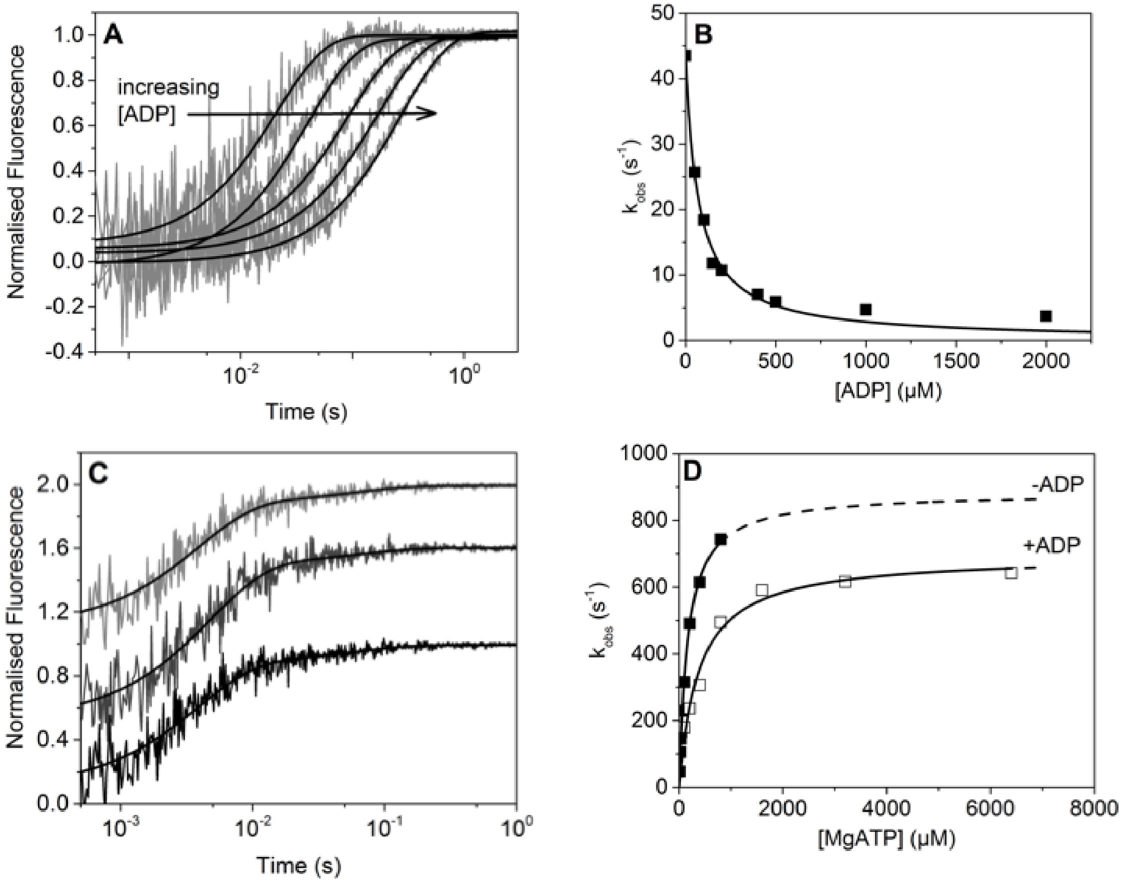

**Figure S2**. ADP affinity for perinatal S1 in the presence of actin. (A) Perinatal acto.S1 was rapidly mixed against a constant concentration of ATP and varied concentrations of ADP. ADP competitively binds to S1 and so slows the dissociation rate as the ADP concentration increases as indicated by the arrow. (B) The value of $((K_D+1)/K_DK_{D^*})$ is the concentration of ADP at which 50% of the S1 is inhibited. When the $k_{obs}$ is plotted against the ADP concentration a $((K_D+1)/K_DK_{D^*})$ value of 69.7 µM is achieved. This was repeated three times and the average values listed in Table S1. (C) Due to the fast rate at which ADP dissociated from S1 in the presence of actin, the dissociation assay was conducted with [ADP] = $((K_{D+1})/K_DK_{D^*})$ (50% of AM with ADP bound). These transient were best described by a double exponential with the $k_{obs}$ of the fast phase increasing with [ATP] while the slow phase was unaffected. The transients have been off-set by 0.25 on the y-axis. (D) Plotting the $k_{obs}$ of the fast phases either with or without ADP present shows how ADP reduces the maximum rate (883.9 ± 21.9 $s^{-1}$ without ADP vs 694.7 ± 28 $s^{-1}$ with ADP). The $K_T$ increases to 377.4 ± 58.4 µM in the presence of ADP compared to 163.9 ± 10.5 µM. i.e. the data are compatible with ADP release being at least 694 $s^{-1.}$

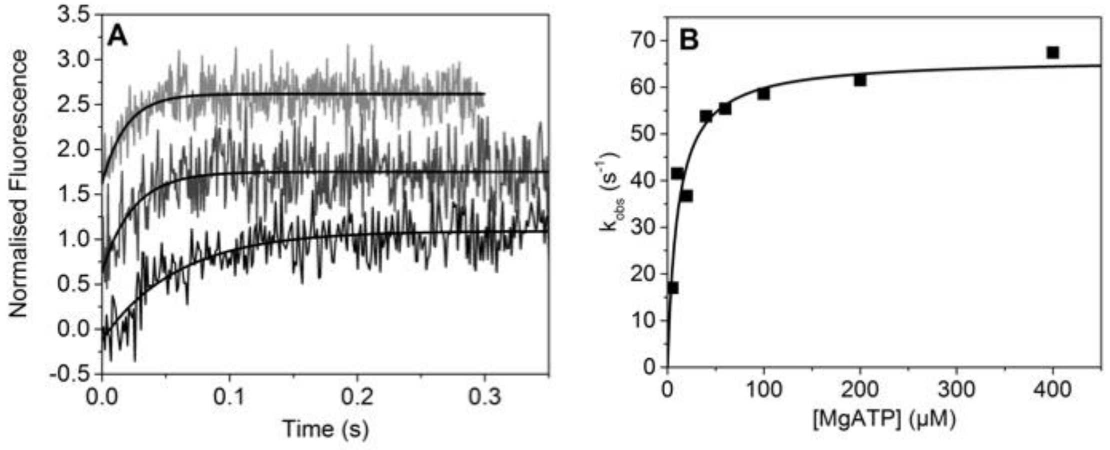

**Figure S3.** Tryptophan fluorescence of ATP binding to perinatal S1. (A) Example transients of the increase in intrinsic tryptophan fluorescence upon the binding of ATP to perinatal S1. This gave a relatively small Δfl (2.5% - 4%) and were best described by a single exponential. (B) When the $k_{obs}$ is plotted against the ATP concentration the data can be best described by a hyperbola which yields a maximum rate $k_{obs} = k_H + k_{-H}$ of 66.1 ± 3.4 $s_{-1}$, the concentration of half-maximum $k_{obs} = K_{50\%}$ of 10.6 ± 2.6 μM, and the second-order rate constant rate constant of 6.2 ± 1.3 $μM^{-1}$ $s^{-1}$.

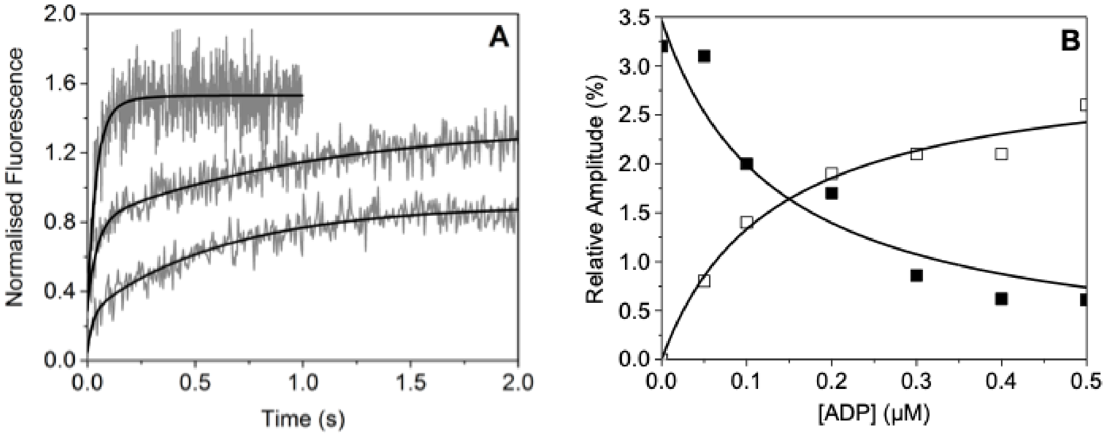

**Figure S4.** ADP affinity for Peri S1 in the absence of actin. (A) S1 was incubated with varied concentrations of ADP (0-0.5 μM after mixing) and rapidly mixed with ATP. In the absence of ADP the transient was best described by a single exponential, whereas when ADP was added the transients could be described by a double exponential. The $k_{obs}$ of these phases did not change as the [ADP] was varied however the fluorescence amplitudes did. (B) Relative amplitudes of the fast phase (closed squares) and slow phase (open squares) when plotted against the [ADP] shows two hyperbolic dependencies. The half-maximum fluorescence change = 0.15 μM and 0.06 μM for the fast and slow phase respectively. This gave an average ADP affinity of 0.105 ±0.045 μM. The $k_{obs}$ of the slow phase represents the ADP release rate constant from S1 = 1.17 ± 0.1 $s^{-1}$. The averages of three repeats are summarised in Table S1.

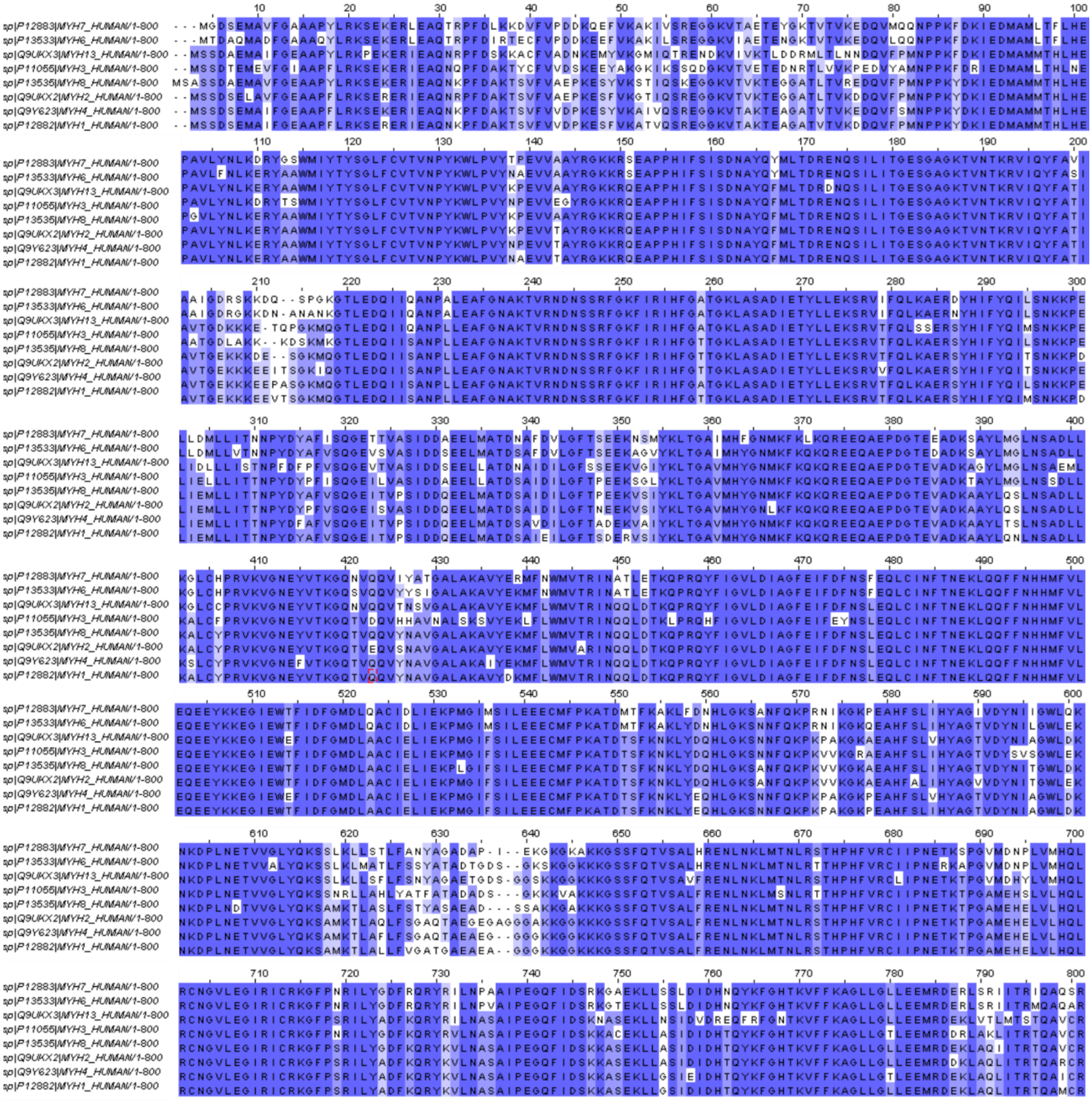

**Figure S5**. Multiple sequence alignment for the 8 myosin class II sequences analysed in this study.
